# Computational Design of Novel Selective Phosphodiesterase 4B Inhibitors from Natural Products: An Integrated Machine Learning and Structure-Based Drug Discovery Approach

**DOI:** 10.64898/2026.05.16.725619

**Authors:** Samson Ayorinde Oni, Moyege Daniel Oyemomi, Osho Adebowale, Abdulfatai Abdulmalik

## Abstract

Selective inhibition of phosphodiesterase 4B (PDE4B) remains a promising strategy for preserving the anti-inflammatory benefit of PDE4 inhibition in chronic obstructive pulmonary disease while reducing PDE4D-associated tolerability liabilities. This study integrated SHAP-interpretable machine learning, natural product virtual screening, hierarchical docking, post-docking MM-GBSA, isoform cross-docking, binding-pocket comparison, ADMET prediction, and 100 ns molecular dynamics simulations to identify PDE4B-selective inhibitors from the LOTUS natural product database. A Random Forest classifier trained on curated ChEMBL PDE4B bioactivity data achieved an external performance with AUC-ROC = 0.955, accuracy = 0.893, F1-score = 0.896, MCC = 0.785, and prioritized 119,698 predicted actives from 276,518 LOTUS compounds. SHAP analysis identified BertzCT and TPSA as major contributors to predicted activity. Sequential Lipinski, PAINS, and QED filtering retained 14,210 candidates for structure-based evaluation. Extra precision docking identified four leads with PDE4B docking scores of -9.123 to -12.080 kcal/mol, all outperforming roflumilast (-7.658 kcal/mol). Cross-docking and post-docking MM-GBSA supported preferential PDE4B binding for three candidates. The top lead, LTS0048837, maintained a stable PDE4B-bound pose during simulation, with comparatively stronger interaction persistence than its PDE4D complex and the roflumilast reference. These findings nominate LTS0048837 as a computationally prioritized PDE4B-selective natural product lead requiring experimental enzyme, cellular, and pharmacokinetic validation.

## 1. Introduction

Globally, chronic obstructive pulmonary disease (COPD) is the fourth major cause of death, with an estimated 3.5 million deaths occurring in 2021 [1]. Global burden estimates show that COPD affects hundreds of millions of individuals worldwide [2]. It is a disease in which airflow becomes increasingly restricted due to chronic inflammation of the airways, remodeling of the small airways, and emphysematous destruction of the parenchyma [3,4]. Despite the availability of bronchodilators and inhaled corticosteroids, current pharmacological therapies primarily reduce symptoms and exacerbation risk rather than consistently reversing or halting the underlying progression of COPD [5,6]. Furthermore, existing anti-inflammatory agents such as roflumilast, while effective in reducing exacerbations in select patients, are limited by dose-dependent adverse effects that restrict their broader clinical application, highlighting the importance of safer and more selective therapeutic approaches [11].

Phosphodiesterase 4 (PDE4) is an enzyme family that breaks down cyclic adenosine monophosphate (cAMP), a key second messenger regulating inflammatory signaling in airway smooth muscle, alveolar macrophages, and neutrophils [7,8]. PDE4 inhibition elevates intracellular cAMP, inhibiting the secretion of pro-inflammatory mediators, including TNF-α, IL-6, and IL-8, and leukotriene B4-cytokines centrally implicated in COPD pathophysiology [5,9]. Roflumilast is the sole approved oral PDE4 inhibitor for severe COPD, and confers modest but clinically meaningful reductions in exacerbation frequency; however, its therapeutic benefit is curtailed by dose-dependent adverse effects including nausea, diarrhea, and weight loss, which limit dose escalation [10,11]. These side effects are mechanistically attributed to PDE4D inhibition, which modulates emetic signaling and gastrointestinal motility, whereas anti-inflammatory efficacy in the lung is principally mediated by PDE4B [12–14]. Accordingly, PDE4B-selective inhibition has emerged as a promising therapeutic strategy, offering the potential to preserve bronchopulmonary anti-inflammatory activity while avoiding PDE4D-mediated tolerability liabilities [12–15].

Achieving PDE4B selectivity over PDE4D presents a formidable challenge: the two isoforms share approximately 80% sequence identity within the catalytic domain, resulting in a highly conserved active site architecture [16]. Nevertheless, subtle isoform-specific differences in residues lining adjacent binding-pocket regions have been reported and may provide opportunities for selective inhibitor design [16,17]. To date, the computational discovery of selective PDE4B inhibitors from natural product chemical space remains largely unexplored despite the well-established pharmacological productivity of natural product scaffolds [18].

The LOTUS database is one of the most comprehensive curated repositories of natural product occurrence data, comprising over 270,000 compounds with detailed taxonomic provenance [19]. Machine learning (ML)-based virtual screening offers a computationally efficient means of rapidly navigating this large chemical space, enabling bioactivity prediction prior to resource-intensive structure-based docking [20]. Interpretability tools like SHapley Additive exPlanations (SHAP) further enable identification of the most important features underlying predictions [21].

For this study, we created and implemented an integrated computational two-stage pipeline comprising SHAP-interpretable ML-based screening, hierarchical molecular docking, Molecular Mechanics/Generalized Born Surface Area (MM-GBSA) free energy calculation, and 100 ns MD simulations to identify novel, selective PDE4B inhibitors from the LOTUS natural product database. Selectivity was assessed through systematic cross-docking and binding free energy comparison against PDE4D, with structural analysis of binding pocket differences providing a mechanistic basis for observed selectivity. This is, to the best of our knowledge, the first study to combine ML-based screening approach with a comprehensive isoform-selective computational profiling pipeline against the LOTUS natural product chemical space for PDE4B inhibitor discovery.

## 2. Materials and Methods

### 2.1. Experimental Design and Computational Workflow

This in silico study was designed to identify selective PDE4B inhibitors from natural product chemical space using a sequential ML-based and structure-based workflow. The prespecified pipeline comprised: curation of PDE4B bioactivity data, descriptor generation and feature selection, machine-learning model development and SHAP interpretation, LOTUS database screening, physicochemical and PAINS filtration, hierarchical docking against PDE4B, top leads cross-docking against PDE4D for isoform selectivity assessment, post-docking MM-GBSA binding free-energy estimation, 100 ns molecular dynamics simulation of the lead complexes, ADMET prediction, and comparative binding-pocket analysis. Figure 1 presents an overview of the LOTUS database screening pipeline.

**Figure 1.**
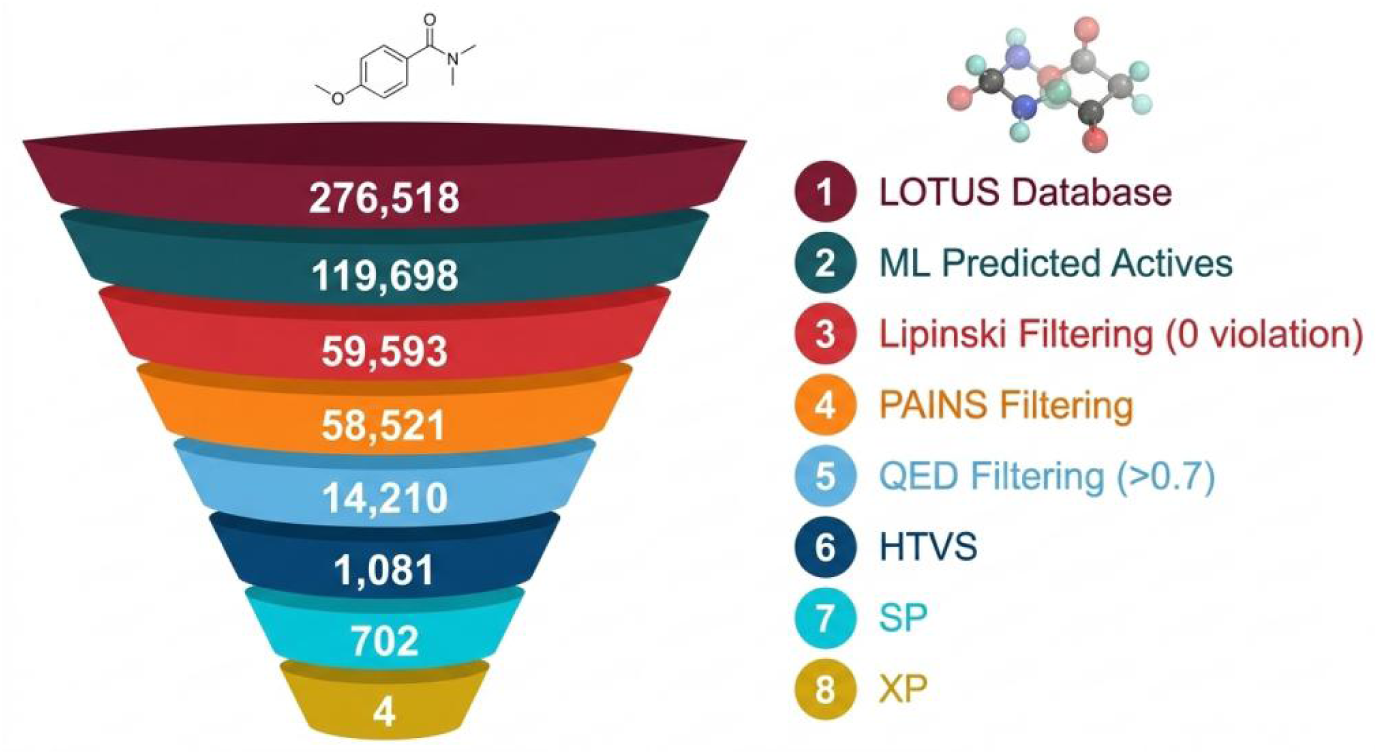
Screening Pipeline.

**Figure 2.**
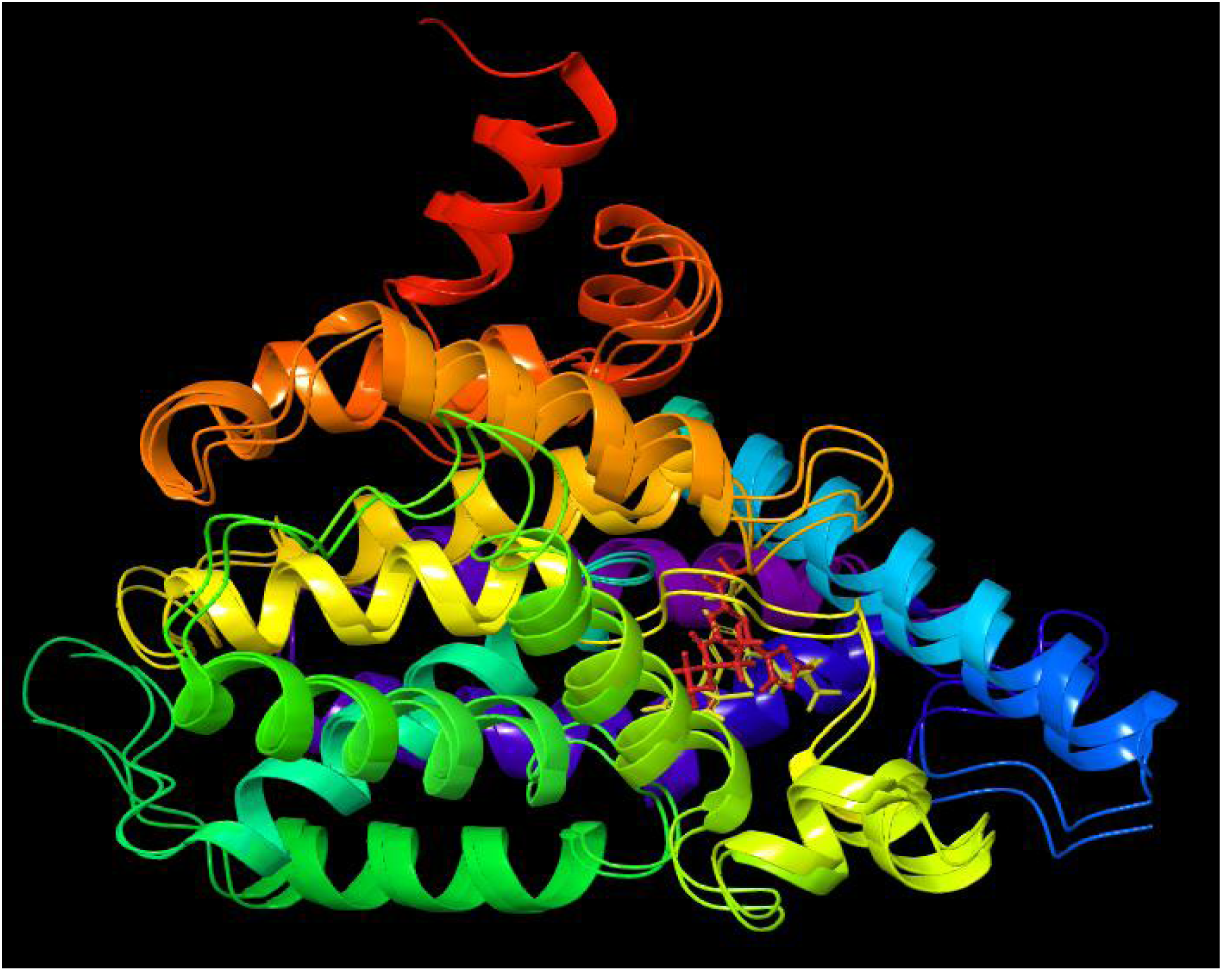
Structural Alignment of PDE4B and PDE4D with their complexes.

### 2.2. Data Collection and Preparation for Machine Learning

Bioactivity data for PDE4B were retrieved from the ChEMBL database [22] using the target identifier, CHEMBL275. A total of 3644 records with experimentally determined IC50 values against human PDE4B were extracted, with activity values represented as pChEMBL Values. After data retrieval and pre-processing (removing duplicate entries, null values, and compounds with invalid SMILES), a dataset of 2,712 compounds was obtained, which comprises 1,325 compounds having pChEMBL Values ≥ 7.0 designated as strong inhibitors and 1387 compounds having pChEMBL Values < 7.0 designated as weak inhibitors (Figure 3). The LOTUS database [19] was downloaded in SMILES format comprising 276,518 natural product structures.

**Figure 3.**
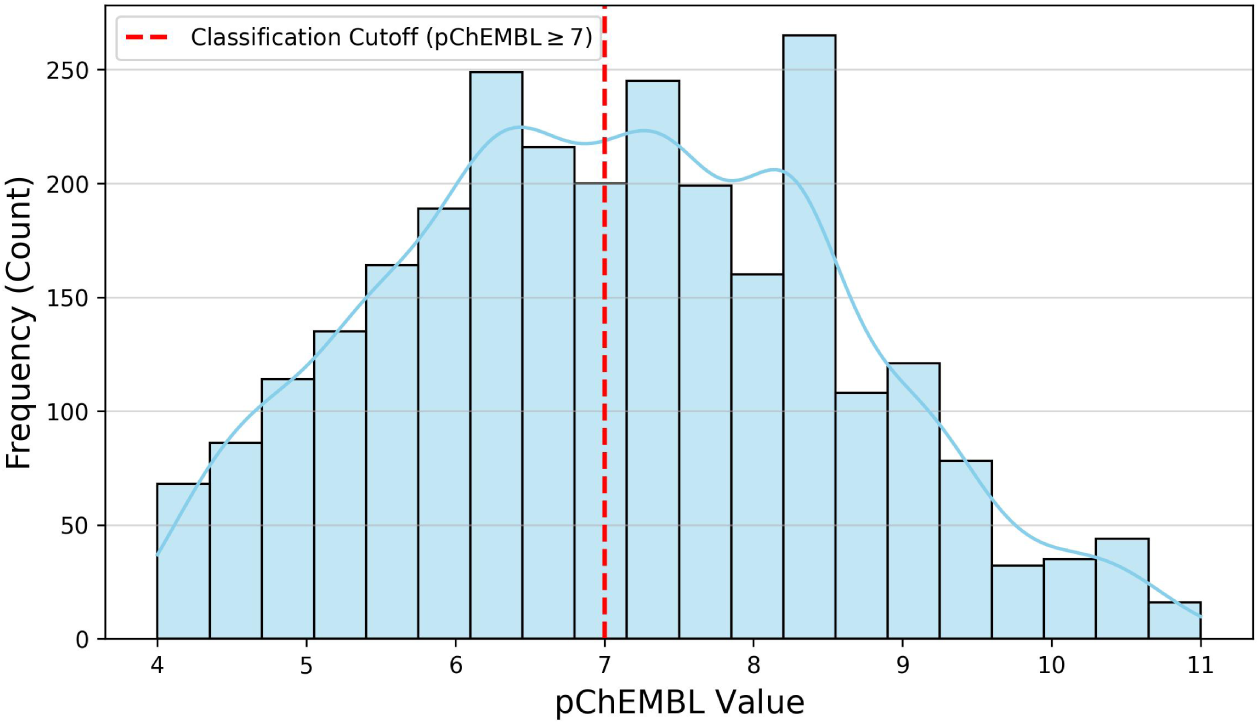
Data distribution of pChEMBL values against PDE4B target.

### 2.3. Molecular Descriptor Calculation and Feature Selection

All compounds in the ChEMBL PDE4B dataset were computed using the Python RDKit library to generate Two-dimensional (2D) molecular descriptors [23], yielding an initial pool of 217 physicochemical and topological descriptors. Feature selection was performed sequentially: (i) low-variance feature removal (threshold = 0.01); (ii) elimination of highly correlated features (Pearson r > 0.90); and (iii) SelectKBest with mutual information scoring retaining the top 20 features [32]. Descriptor values were normalized using StandardScaler.

### 2.4. Machine Learning Model Development and Validation

The curated set of data was divided into a training set (80%) and a held-out test set (20%), using stratified random splitting (random_state = 42). Five classification algorithms were evaluated: XGBoost, RandomForest, SVM_Linear, LightGBM and Gradient Boosting. Hyperparameter optimization was conducted using GridSearchCV with five-fold stratified cross-validation (StratifiedKFold, n_splits = 5, F1-score optimization). The accuracy and Area under the Receiver Operating Characteristic (AUC-ROC) were used to evaluate the performance of the model, F1-score, precision, recall, and Matthew’s Correlation Coefficient (MCC) on the external test set, alongside mean cross-validation F1-score on training partitions. The model that performed best was chosen based on MCC. Model interpretability was quantified using SHAP TreeExplainer, generating global feature importance rankings and beeswarm summary plots (Figure 7).

### 2.5. LOTUS Database Screening and Compound Filtration

The 20 selected training descriptors were computed for all LOTUS compounds using the same RDKit functions employed during model building. Critically, raw descriptor values were transformed using the fitted StandardScaler prior to prediction to ensure feature scale consistency with the training distribution. Predicted strong inhibitors (probability ≥ 0.50) were retained and subjected to the following sequential filters with zero violations tolerated: (i) Lipinski Rule-of-Five: MW ≤ 500 Da, LogP ≤ 5, HBD ≤ 5, HBA ≤ 10, rotatable bonds ≤ 10 [24]; (ii) PAINS substructure filtering via RDKit FilterCatalog [25]; and (iii) QED ≥ 0.70 [26]. An applicability domain assessment was performed using Principal Component Analysis (PCA) projection, overlaying the training set and predicted active LOTUS compounds to evaluate chemical space overlap.

### 2.6. Protein Structure Retrieval and Preparation

Crystal structures of PDE4B (PDB ID: 4KP6; resolution: 1.50Å) and PDE4D (PDB ID: 3G4L; resolution: 2.50Å) were retrieved from the RCSB Protein Data Bank [27]. The preparation of both structures was carried out with the Protein Preparation Wizard in Schrödinger Maestro [28], with addition of hydrogen atoms, bond order assignment, protonation state optimization at pH 7.0 ± 1.0 (PROPKA), water removal beyond 5.0 Å, side chain and loop completion, and restrained energy minimization with OPLS4.

### 2.7. Docking Protocol Validation and Receptor Grid Generation

The Receptor Grid Generation Module in Maestro was used to create the glide receptor grids, which focused on the binding site of each co-crystallized ligand target. Grid box coordinates for 4KP6 receptor were x= -38.28, y=59.7 and z=112.05; for 3G4L, the grid box coordinates were x= -38.99, y=60.46 and z=110.1. The re-docked extra precision (XP) mode of the native ligands of 4KP6 and 3G4L were used to validate the docking protocol.

### 2.8. Hierarchical Molecular Docking

Predicted strong inhibitors that passed the filtering steps (as described in section 2.5), were built in Maestro with LigPrep module for docking (OPLS4 force field, pH 7.0 using Epik). Roflumilast was prepared identically as the reference compound. All the compounds were docked against 4KP6 in three sequential stages: High Throughput Virtual Screening (HTVS), Standard Precision (SP), and eXtra Precision (XP) docking processes. The top 1081 compounds (docking scores ranging -7.00kcal to -9.79kcal) from the initial HTVS docking advanced to SP docking. The final screening was performed with XP docking for the top 702 compounds from the SP docking (docking scores ranging from -7.00kcal to -10.49kcal). The top four XP-ranked compounds were cross-docked against 3G4L using the same XP protocol. Roflumilast was docked against PDE4B and PDE4D using XP protocol. Two-dimensional protein-ligand interaction diagrams of the top 4 compounds and Roflumilast complexed with 4KP6 and also with 3G4L were generated in Maestro.

### 2.9. Molecular Mechanics/Generalized Born Surface Area (MMGBSA) Binding Free Energy Calculations

MM-GBSA analysis estimates the ligand-protein binding free energy and provides a refined assessment of the predicted binding strength beyond docking scores alone [30]. Binding free energies of the top compounds against 4KP6 and 3G4L were computed using Prime MM-GBSA (OPLS4 force field, VSGB 2.0 solvation model) on XP docked poses [29,30]. These calculations were performed as post-docking single-pose MM-GBSA estimates.

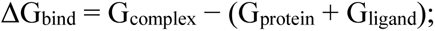

where G = molecular mechanics energy + solvation free energy.

### 2.10. Molecular Dynamics Simulations

Molecular dynamics simulations of 100 ns were carried out for three complexes: PDE4B-LTS0048837, PDE4D-LTS0048837, and PDE4B-roflumilast, using Desmond (Schrödinger) with the OPLS4 force field. The solvation of each system was added in a TIP4P orthorhombic water box of size 10Å. Na+/Cl- ions were added to neutralize the system and maintain 0.15 M NaCl [30]. After the minimization of energy, production MD was run under NPT conditions (T = 310 K, Nose-Hoover thermostat; P = 1.01325 bar, Martyna-Tobias-Klein barostat). Post-simulation analyses included protein backbone RMSD, ligand RMSD after fitting on the protein backbone (Lig Fit Prot), per-residue Cα RMSF, protein-ligand contact histograms, 2D simulation interaction diagrams, and contact timeline plots.

### 2.11. ADMET and Physicochemical Property Prediction

The top four compounds and roflumilast were analyzed for their predicted ADMET and physicochemical properties using SwissADME [31]. Selected parameters included MW, iLogP, TPSA, HBD, HBA, rotatable bonds, GI absorption, BBB permeability, likelihood of P-gp substrate, CYP inhibition profiles (CYP1A2, 2C19, 2C9, 2D6, 3A4), and Bioavailability Score.

### 2.12. Binding Pocket Comparison and Structural Basis of Selectivity

PDE4B (4KP6) and PDE4D (3G4L) structures were structurally aligned in Maestro. Binding pocket residues within 5 Å of the top docked compound were identified in both targets; conserved and non-conserved residues were catalogued and their interactions with the top compound analyzed. The non-conserved residues provided a structural rationale for observed selectivity.

## 3. Results

### 3.1. Machine Learning Model Performance

The five ML models were evaluated; performance metrics are summarized in Table 1. The RandomForest classifier achieved the highest discriminatory performance on the external test set: AUC-ROC = 0.955, accuracy = 0.891, F1-score = 0.891, precision = 0.913, recall = 0.871, and MCC = 0.784. The mean cross-validation F1-score on training partitions was 0.9847. Confusion matrix result of the best model (Figure 4) shows that the Random Forest classifier correctly identified 242 strong inhibitors as strong inhibitors (TP) and 242 weak inhibitors as weak inhibitors (TN), while 23 weak inhibitors were misclassified as strong inhibitors (FP) and 36 strong inhibitors were misclassified as weak inhibitors (FN). The ROC curves are shown in Figure 5, with the Random Forest model having the best discriminatory performance, achieving the highest AUC-ROC value of 0.955, indicating strong ability to distinguish between weak and strong inhibitors [32]. The predicted probability distribution showed limited classification ambiguity (Figure 6). Most weak inhibitors clustered at low probabilities (<0.20), while most strong inhibitors clustered at high probabilities (>0.80). Only few compounds appeared around the 0.50 decision threshold.

**Figure 4.**
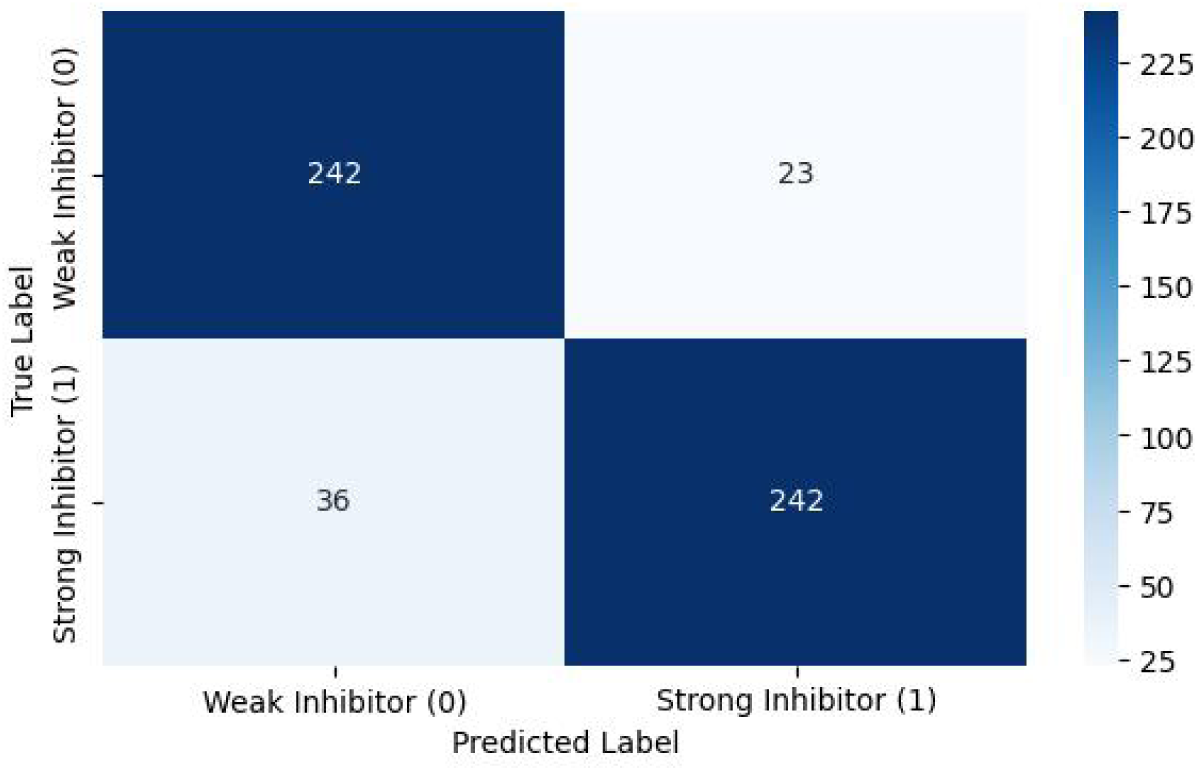
Confusion Matrix of the Random Forest Model.

**Figure 5.**
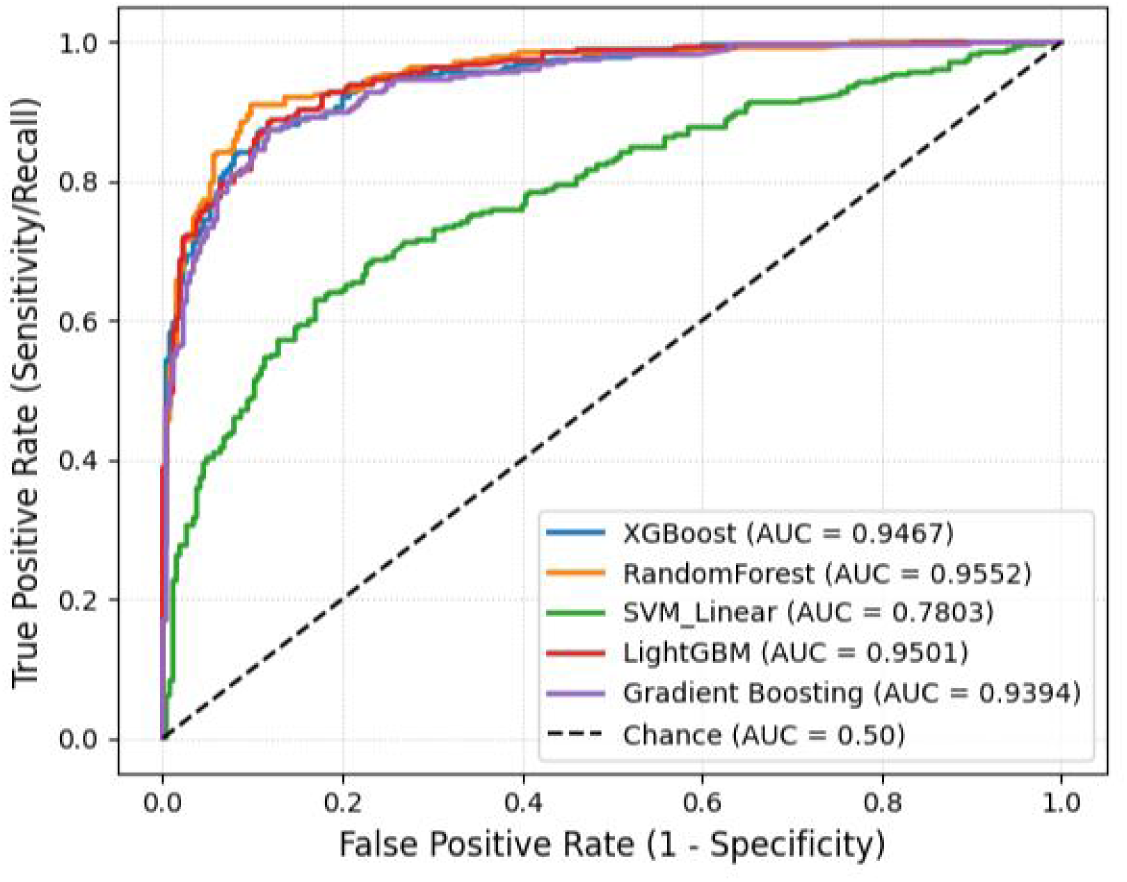
Receiver Operating Characteristics (ROC) Curve Comparison Among the Five Models.

**Figure 6.**
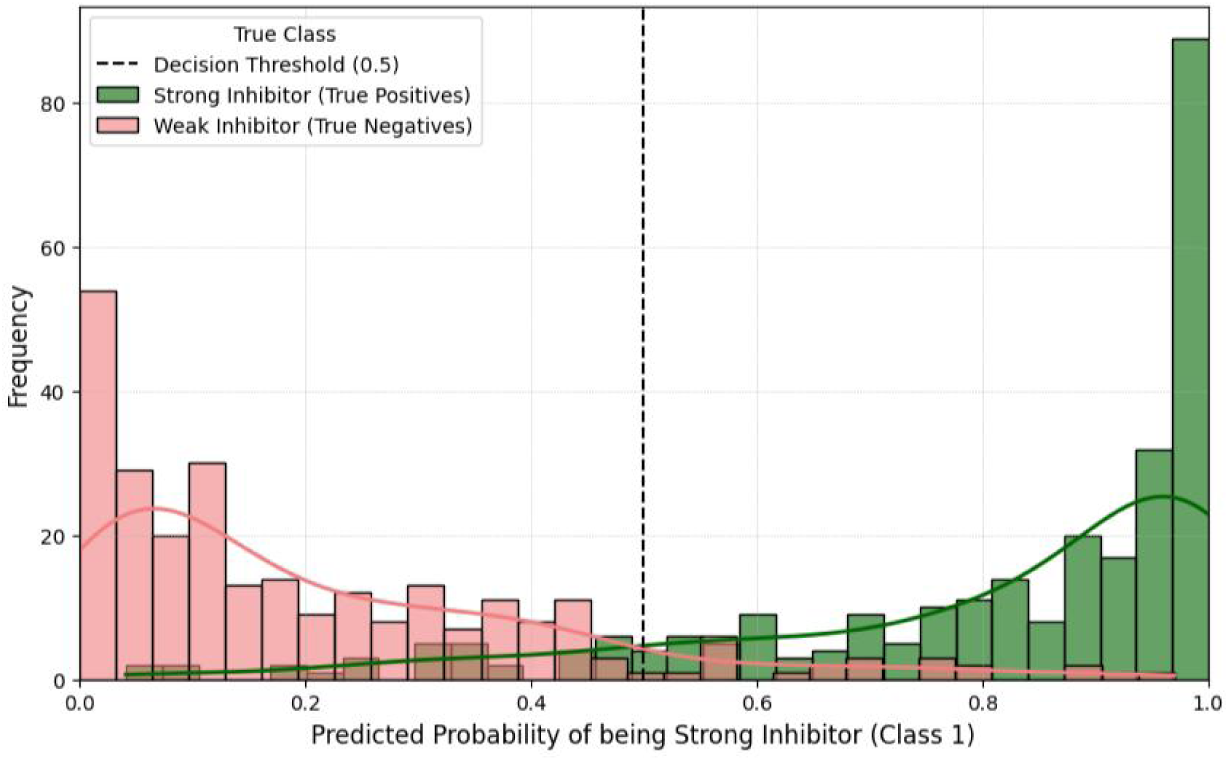
Predicted Probability Distribution (Random Forest Model)

**Figure 7.**
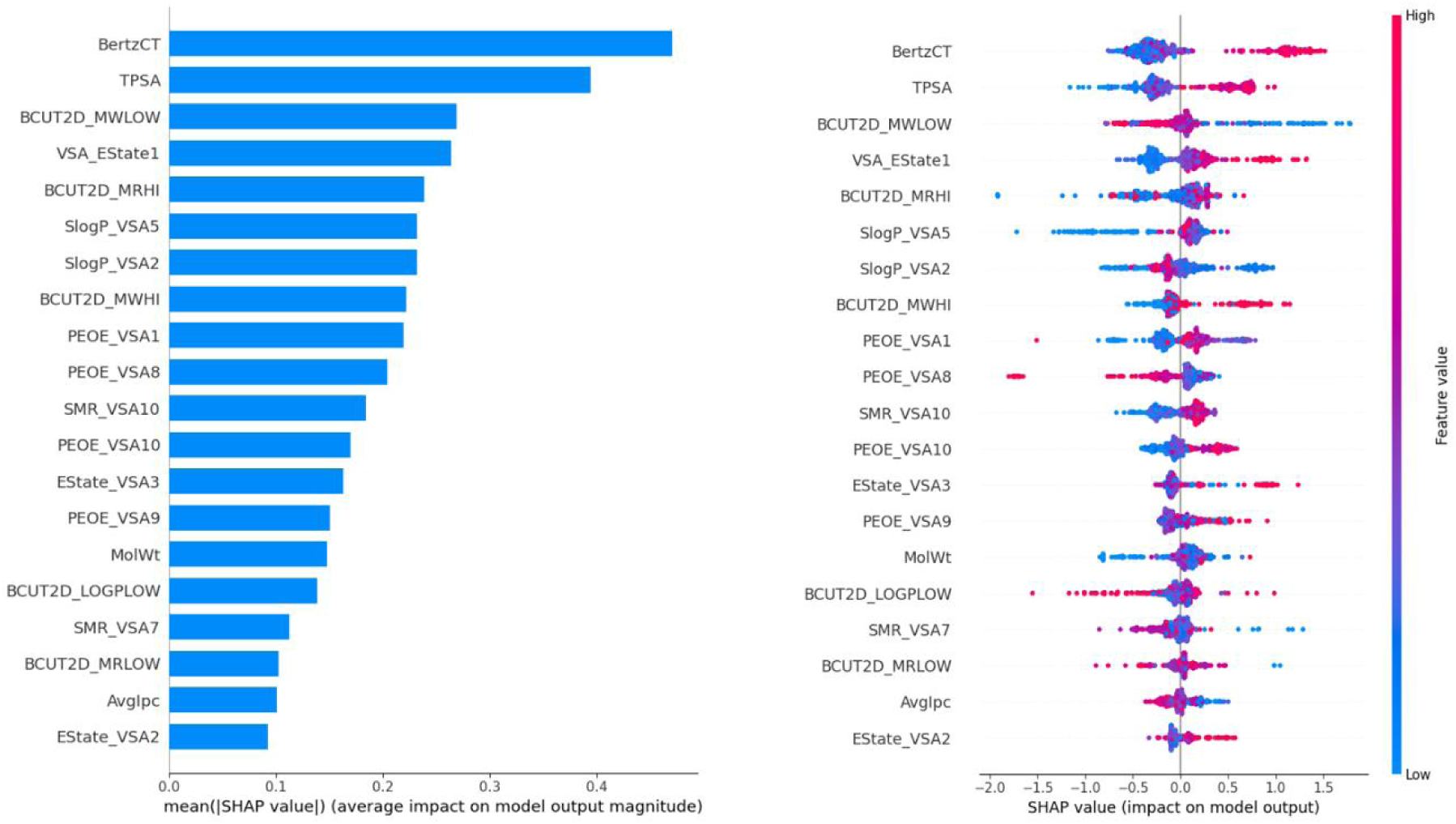
SHAP Analysis.

**Table 1.**
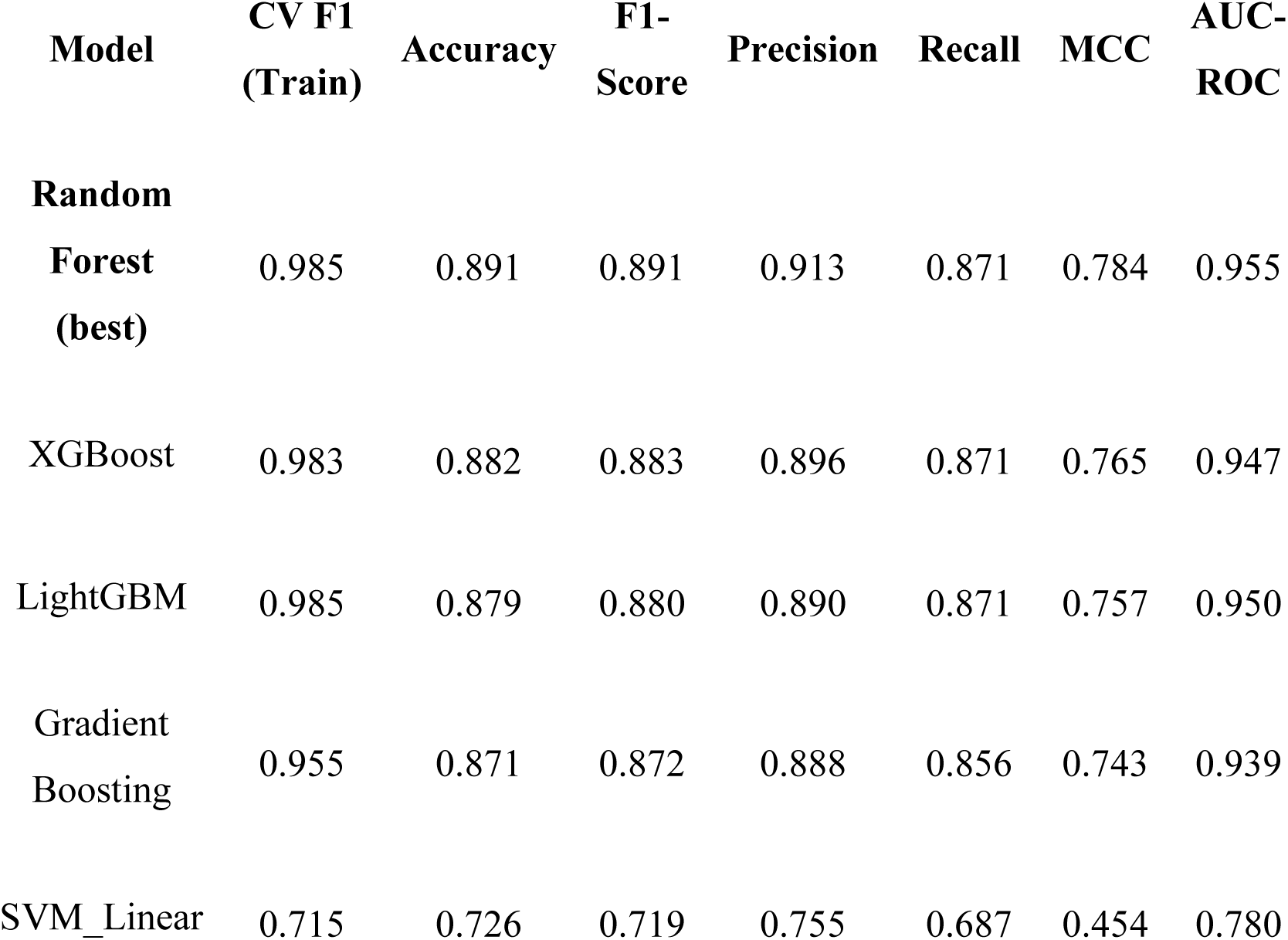
Comparative Performance Metrics of all ML Classification Models on the External Test Set.

SHAP analysis (Figure 7) identified BertzCT, TPSA, and BCUT2D_MWLOW as the three most influential descriptors contributing to Random Forest predictions. The SHAP bar plot ranked these descriptors highest based on mean absolute SHAP values, indicating that they had the highest mean contribution to the model’s output. The beeswarm plot further suggested that higher BertzCT and TPSA values generally shifted predictions toward the strong-inhibitor class, whereas lower BCUT2D_MWLOW values appeared to contribute positively to strong-inhibitor prediction. Overall, the SHAP output suggest that the model distinguished weak from strong inhibitors using a combination of molecular complexity, polarity-related surface properties, and atom-distribution descriptors, which are chemically plausible determinants of ligand activity.

### 3.2. LOTUS Screening Funnel and Compound Filtration

Application of the ML model to the 276,518 LOTUS compounds predicted 119,698 compounds as strong PDE4B inhibitors (probability ≥ 0.50). Sequential filtration removed 60,105 compounds for at least one Lipinski violations, 1,072 compounds as PAINS, and 44,311 compounds with QED < 0.70, yielding 14,210 compounds for structure-based screening. PCA applicability domain analysis showed substantial overlap between the predicted LOTUS actives and the training-set chemical space, indicating that a large proportion of the predicted actives fall within or near the model’s applicability domain (Figure 8).

**Figure 8.**
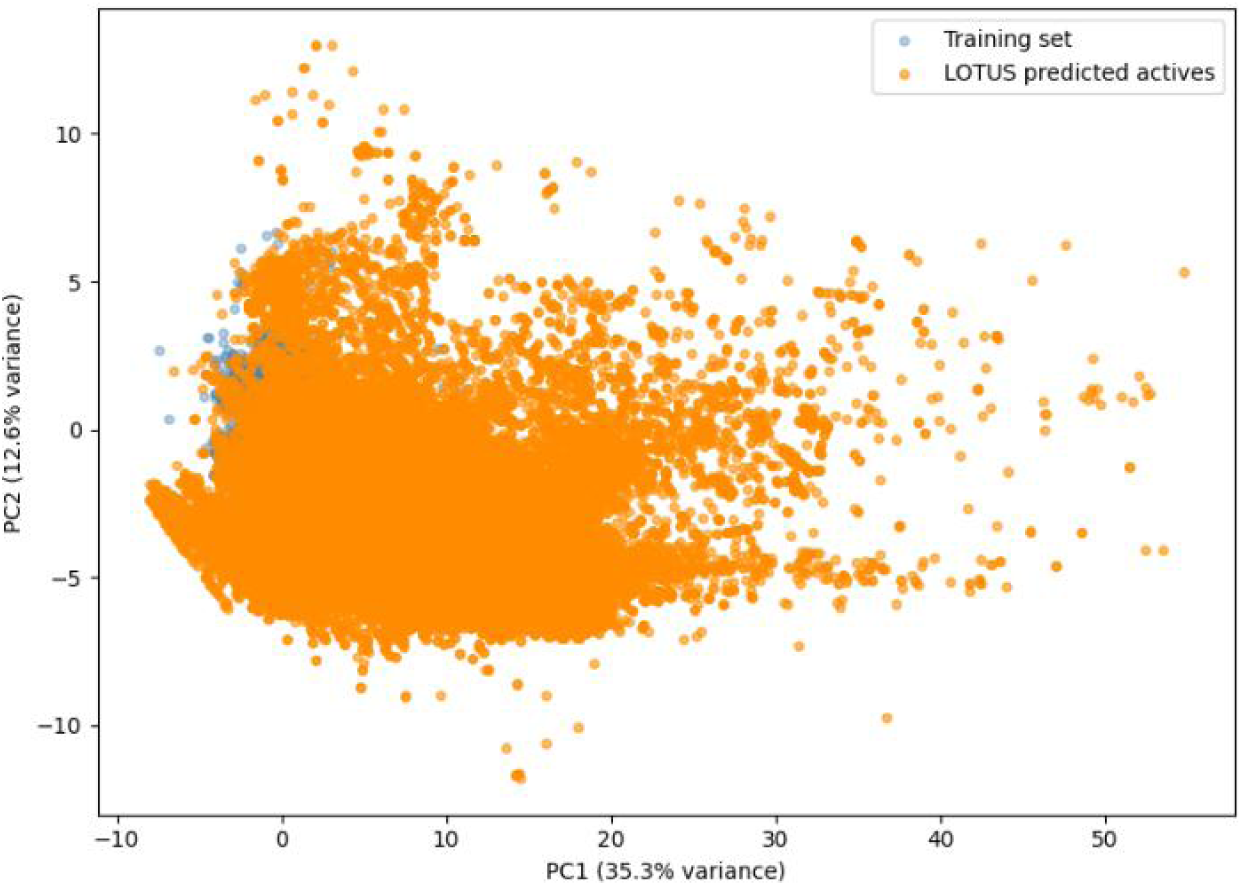
PCA Applicability Domain Analysis.

### 3.3. Docking Validation and PDE4B Docking Results

Re-docking of the native 4KP6 and 3G4L ligands reproduced their crystallographic poses with RMSD of 1.37Å and 1.53Å for 4KP6 and 3G4L, respectively (accepted threshold: < 2.0 Å) [33], confirming docking reliability. Ligand preparation generated 18,383 prepared ligand structures from the 14,210 predicted active compounds. The additional 4,173 structures arose from the generation of alternative ligand states during LigPrep, including possible protonation/ionization states, tautomeric forms, stereoisomeric variants, and energy-minimized 3D conformations. Hierarchical screening advanced the 18,383 compounds through HTVS, 1081 through SP, and 702 through XP docking. The top four lead compounds (Table 2) exhibited docking scores of -9.123 to -12.080 kcal/mol against 4KP6, all more negative than roflumilast -7.658kcal/mol.

**Table 2.**
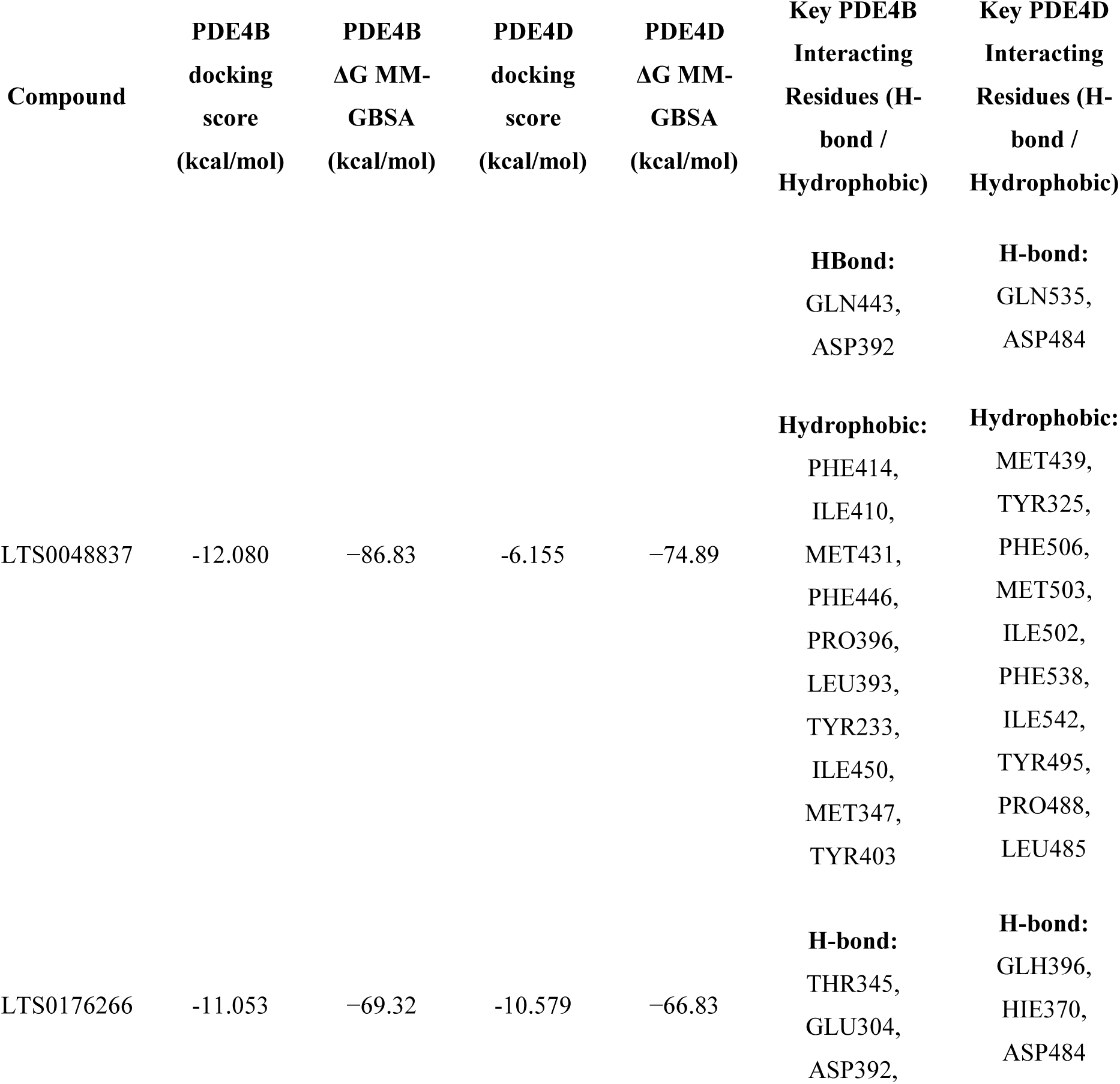
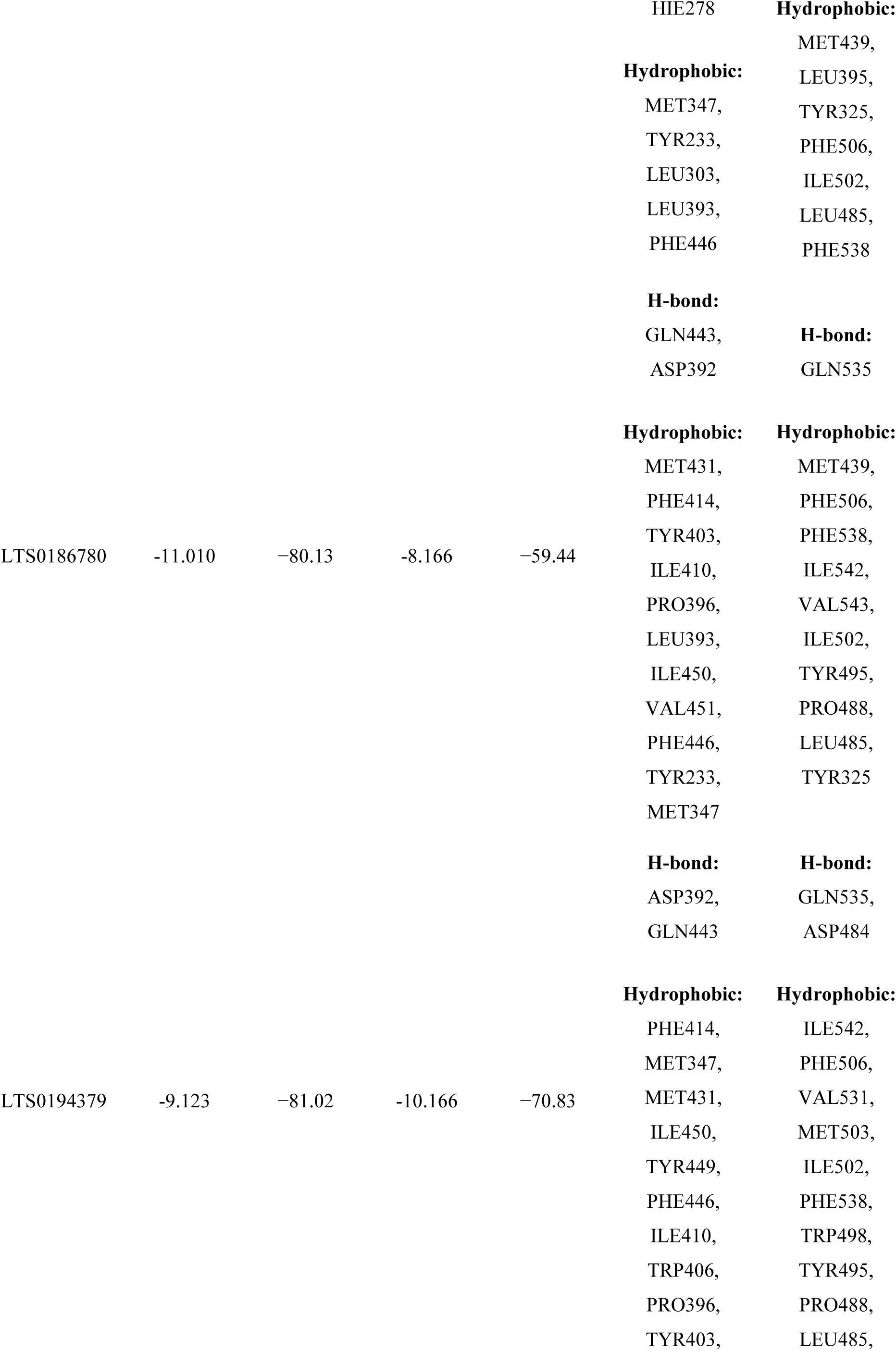
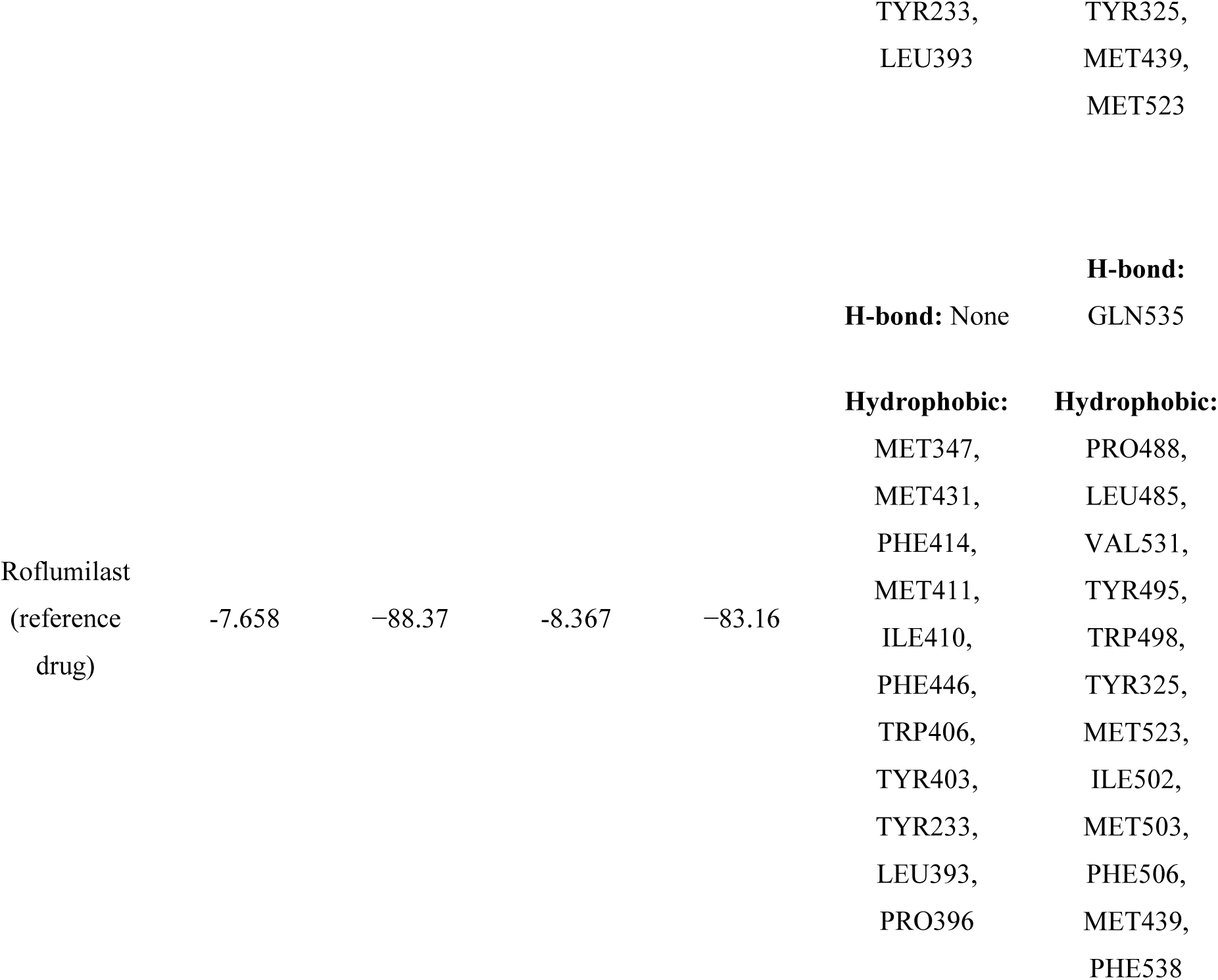
XP Docking Score and MM-GBSA Binding Free Energies of the Top Four Lead Compounds and Roflumilast against PDE4B (4KP6) and PDE4D (3G4L)

The top compound, LTS0048837, demonstrated a docking score of -12.080 kcal/mol and formed 3 hydrogen bonds with active site residues ASP392 and GLN443. These interactions are analogous to interaction patterns reported by Cai et al. [17]. Interaction diagrams for all the four lead compounds and roflumilast are shown in Figures 9 (a-e) and 10 (a-e)

**Figure 9.**
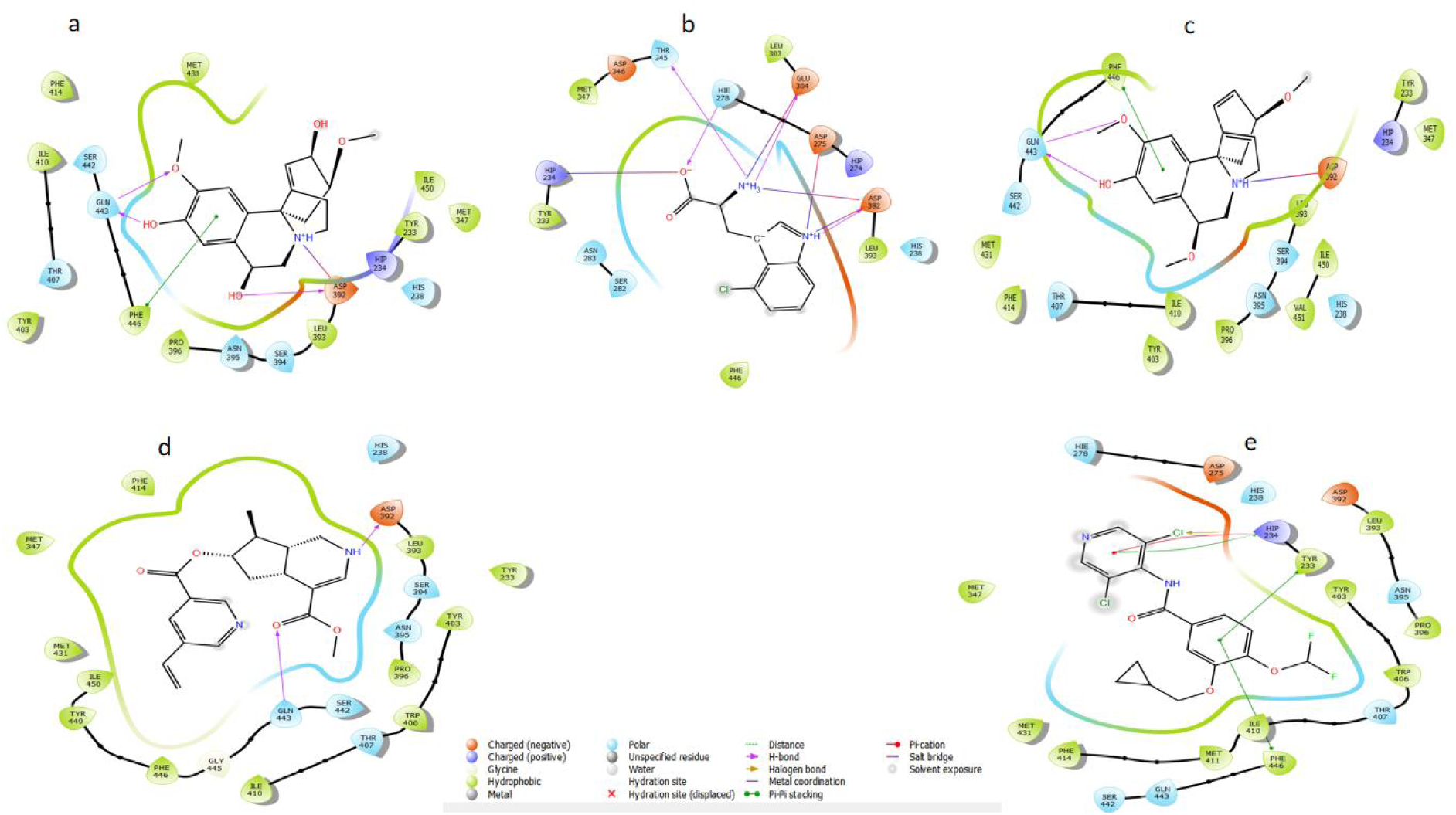
Two-dimensional protein-ligand interaction diagrams of the top four lead compounds and roflumilast bound to PDE4B (4KP6) *a. Two-dimensional protein-ligand interaction of 4KP6-LTS0048837 complex; b. Two-dimensional protein-ligand interaction of 4KP6-LTS0176266 complex; c. Two-dimensional protein-ligand interaction of 4KP6-LTS0186780 complex; d. Two-dimensional protein-ligand interaction of 4KP6-LTS0194379 complex; e. Two-dimensional protein-ligand interaction of 4KP6-Roflumilast complex*

**Figure 10.**
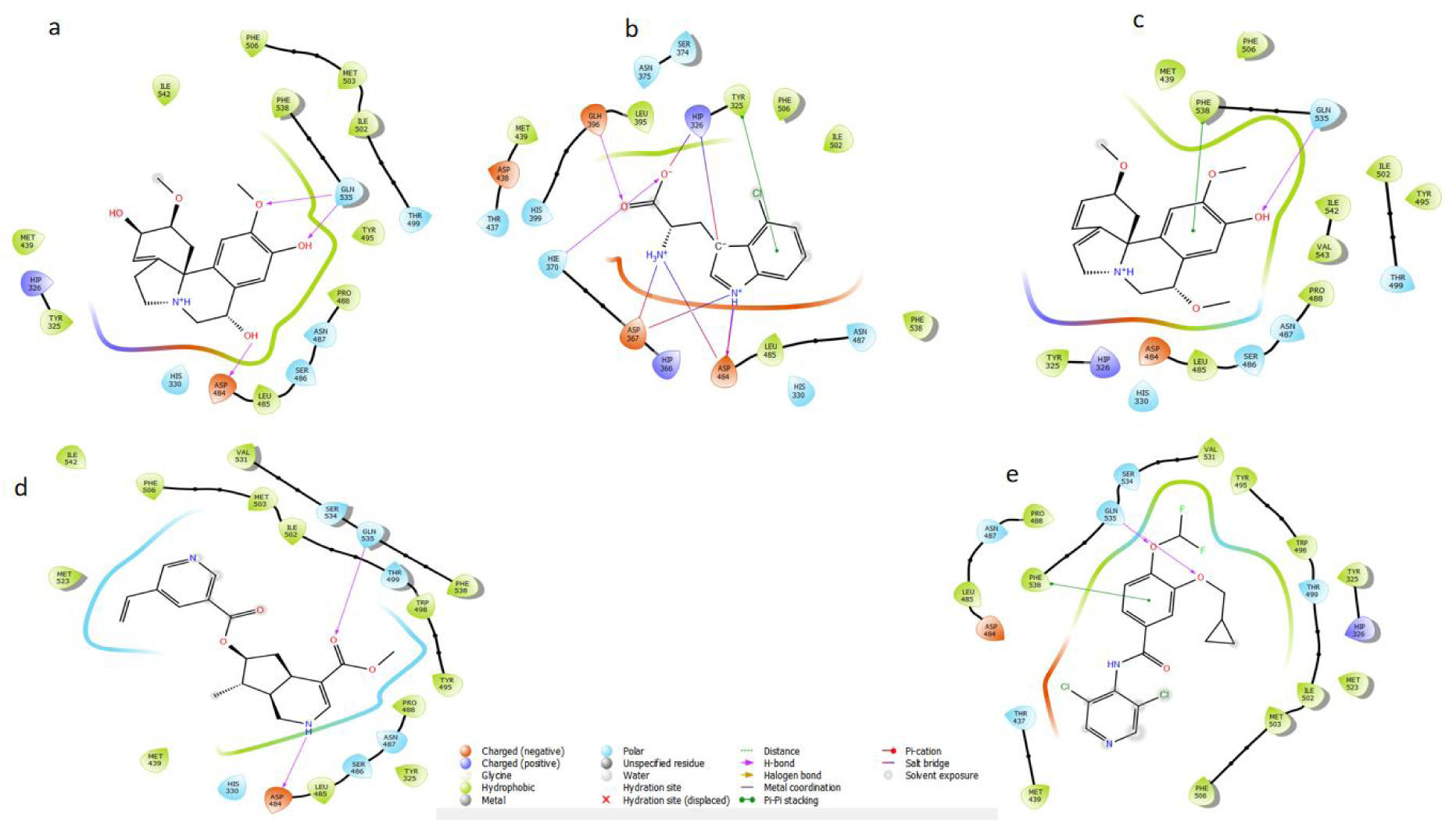
Two-dimensional protein-ligand interaction diagrams of the top four lead compounds and roflumilast bound to PDE4D (3G4L) *a. Two-dimensional protein-ligand interaction of 3G4L-LTS0048837 complex; b. Two-dimensional protein-ligand interaction of 3G4L-LTS0176266 complex; c. Two-dimensional protein-ligand interaction of 3G4L-LTS0186780 complex; d. Two-dimensional protein-ligand interaction of 3G4L-LTS0194379 complex; e. Two-dimensional protein-ligand interaction of 3G4L-Roflumilast complex*

**Figure 11.**
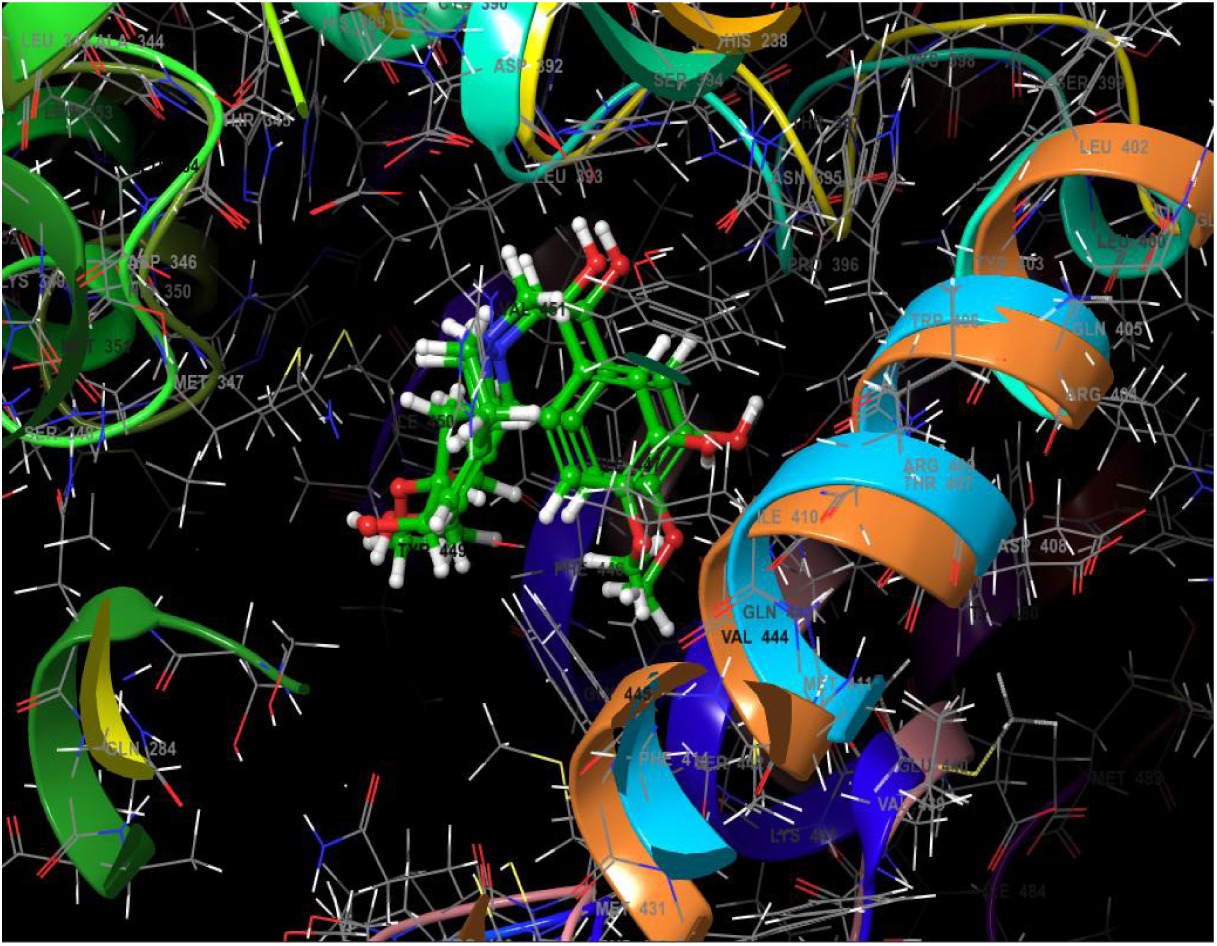
Structural alignment of PDE4B (4KP6) and PDE4D (3G4L) showing LTS0048837 binding.

**Figure 12.**
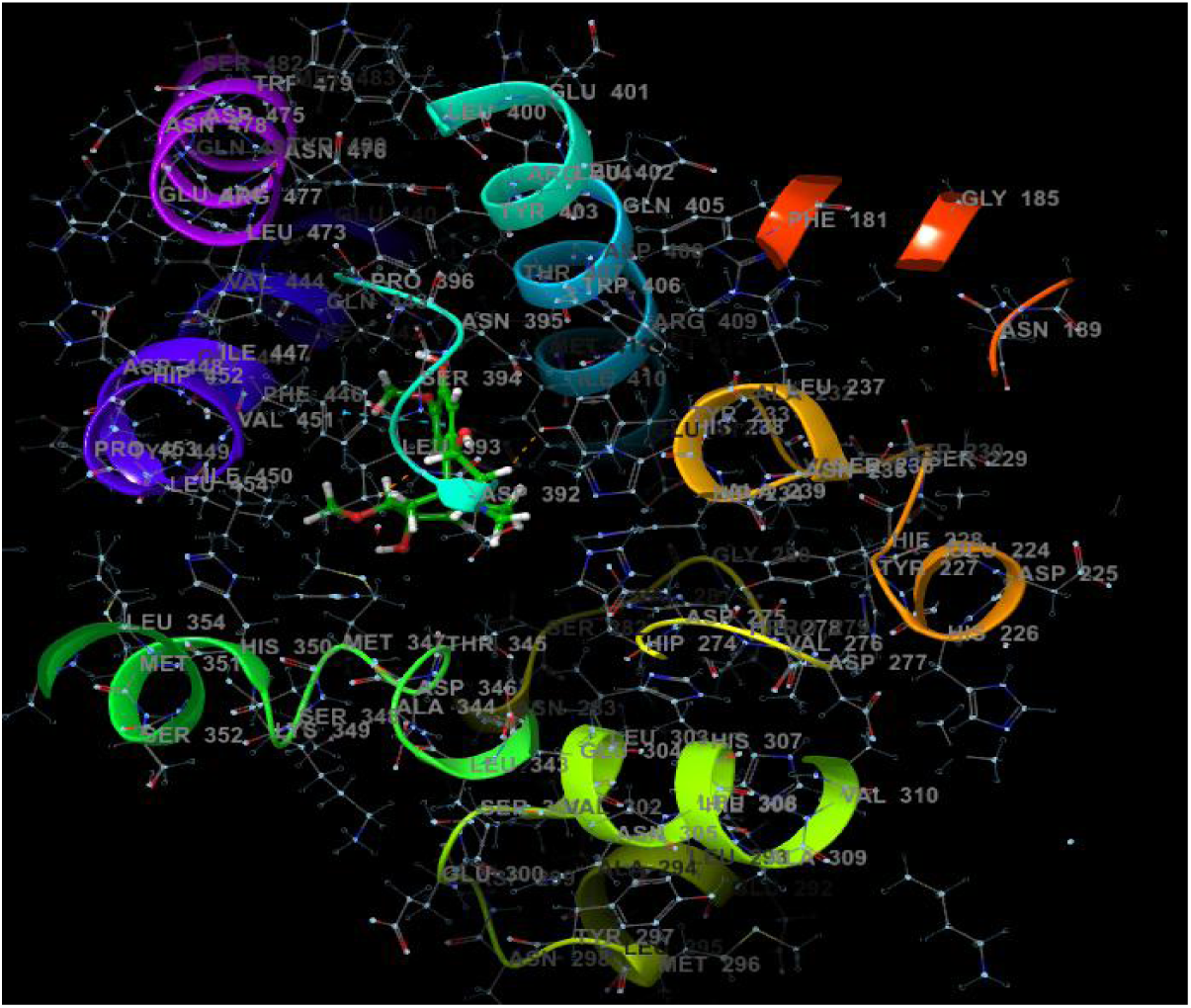
PDE4B (4KP6) binding pocket visualization.

**Figure 13.**
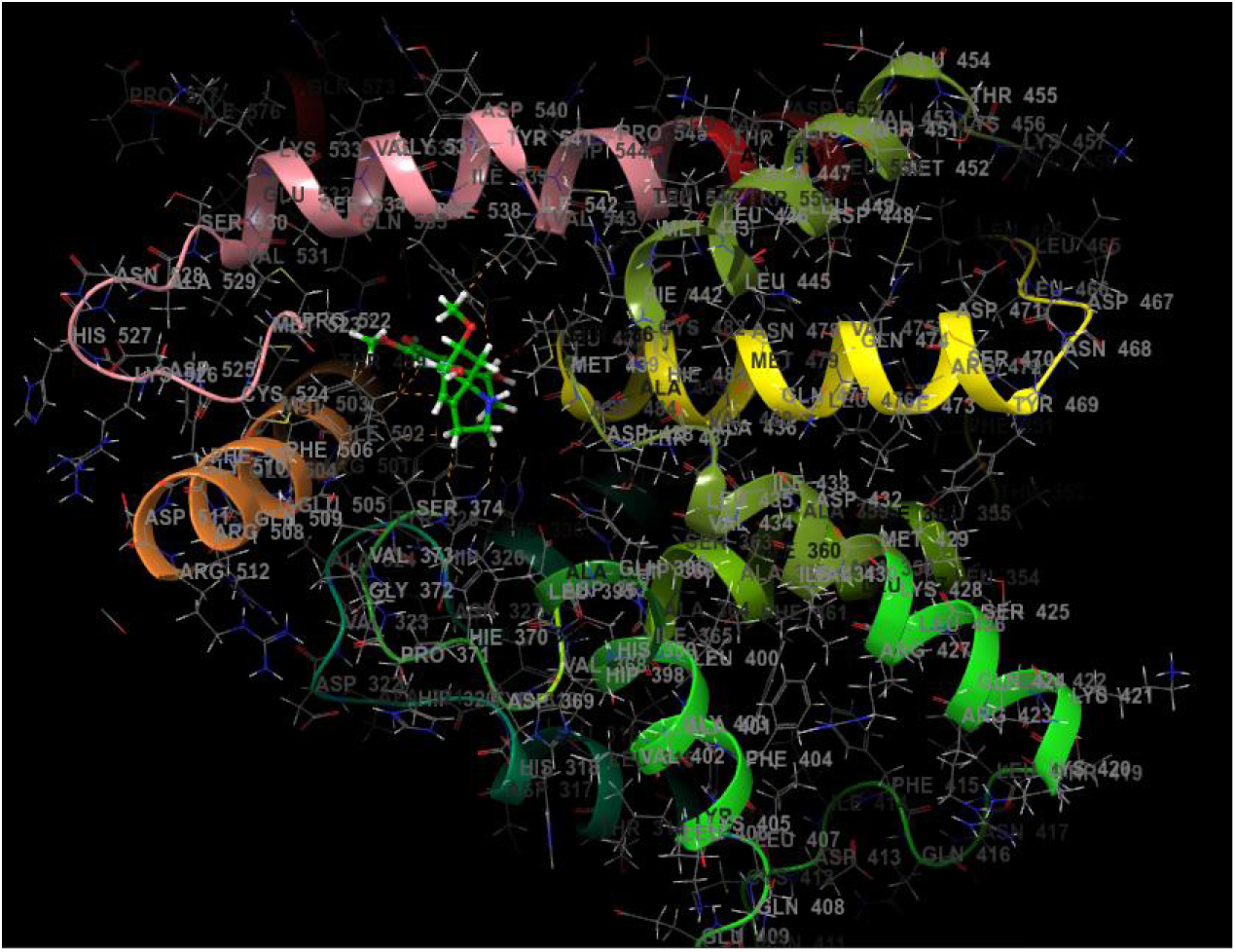
PDE4D (3G4L) binding pocket visualization.

### 3.4. Selectivity Profiling: PDE4B vs. PDE4D

Cross-docking of the four PDE4B leads against PDE4D consistently yielded less favorable docking scores and, critically, the binding free energies (MM-GBSA) for the four compounds were less negative for PDE4D than for PDE4B, except for LTS0194379 which had a better docking score against PDE4D (Table 2). The top compound, LTS0048837, showed MM-GBSA score margin (ΔΔG MM-GBSA) of 11.94 in favor of PDE4B, the largest observed in the series.

Structural alignment of PDE4B (4KP6) and PDE4D (3G4L) identified six binding pocket residues within 5 Å of the top docked compound in each target. The PDE4B pocket comprises Tyr233, Met347, Leu393, Ile410, Phe414, and Phe446, while the corresponding PDE4D pocket is lined by Met439, Leu485, Ile502, Phe506, Phe538, and Ile542. The interaction analysis showed that LTS0048837 engages a hydrophobic PDE4B pocket, with Tyr233 contributing a π–π stacking interaction rather than a hydrogen bond (Figures 9a and 17a). In the MD-derived 2D interaction profile (Figure 17a), Tyr233 maintained contact with LTS0048837 for approximately 47% of the simulation time. Therefore, Tyr233 may contribute to ligand stabilization through aromatic stacking within the PDE4B pocket, while ASP392 and GLN443 provide the major polar interaction anchors.

### 3.5. Molecular Dynamics Simulations

#### 3.5.1. Root Mean Square Deviation (RMSD) and System Equilibration

Protein backbone and ligand RMSD profiles over 100 ns are shown in Figure 14(a-c). For the 4KP6-LTS0048837 complex, the protein backbone RMSD increased during the initial relaxation phase and then remained largely within approximately 1.5-2.4 Å for the rest of the trajectory, with an average value of 1.88 ± 0.25 Å after equilibration. The ligand RMSD after fitting on the protein backbone fluctuated around a stable band of approximately 2.0-3.2 Å (2.62 ± 0.34 Å), indicating that LTS0048837 retained its binding-site orientation without major displacement from the PDE4B pocket. In contrast, the 3G4L-LTS0048837 complex showed a progressive increase in protein RMSD, especially after ∼65 ns, approaching approximately 2.8-3.1 Å towards the end of the simulation. Although the ligand RMSD in the PDE4D complex remained mostly around 1.0-2.0 Å, the increasing protein backbone deviation indicates greater conformational adjustment of PDE4D around the bound ligand. The 4KP6-roflumilast reference complex showed a protein RMSD generally around 1.8-2.6 Å, but its ligand RMSD displayed broader fluctuations, including excursions above 3.5 Å, suggesting greater ligand mobility than LTS0048837 in the PDE4B active site.

**Figure 14.**
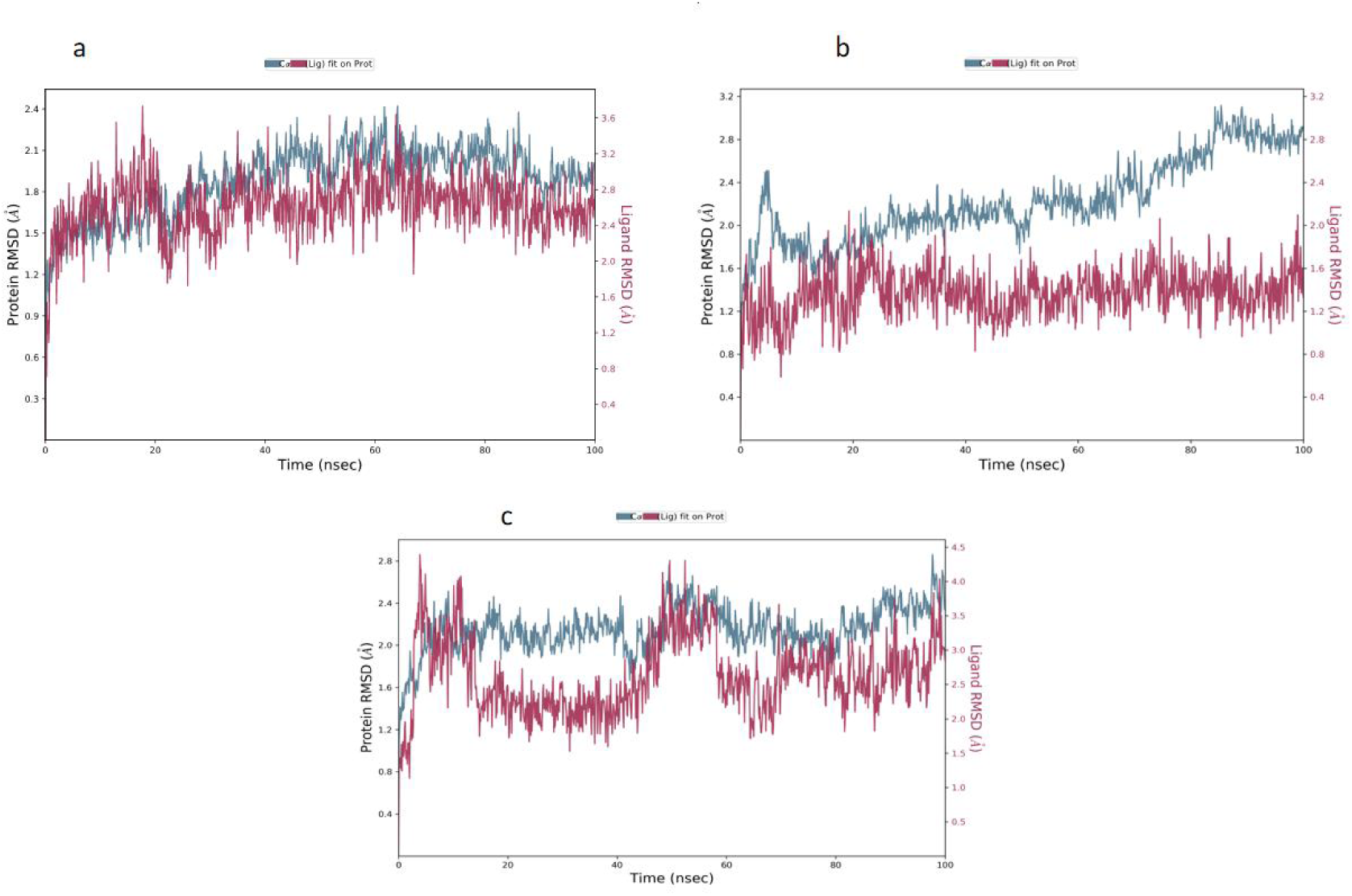
(a-c). 100 ns MD Simulation RMSD plots. a. Root Mean Square Deviation (RMSD) plot for the 100 ns MD simulation of the 4KP6-LTS0048837 complex; b. RMSD plot for the 100 ns MD simulation of the 3G4L-LTS0048837 complex; c. RMSD plot for the 100 ns MD simulation of the 4KP6-roflumilast complex.

#### 3.5.2. Root Mean Square Fluctuation (RMSF) Analysis

Per-residue Cα RMSF profiles are presented in Figure 15(a-c). The 4KP6-LTS0048837 complex showed generally low residue flexibility across most of the protein, with an average RMSF of ∼1.10 Å. The main fluctuations were concentrated in flexible loop/terminal regions, including a peak around residues 130-140, a second region around residues 260-275, and the terminal region near the end of the sequence. The 3G4L-LTS0048837 complex displayed a similar low-fluctuation baseline across much of the protein but showed more pronounced localized peaks around residues 260-280 and at the C-terminal region, where RMSF approached approximately 4-5 Å. The 4KP6-roflumilast complex also showed low-to-moderate residue fluctuation across most residues, but with terminal and loop-region peaks.

**Figure 15.**
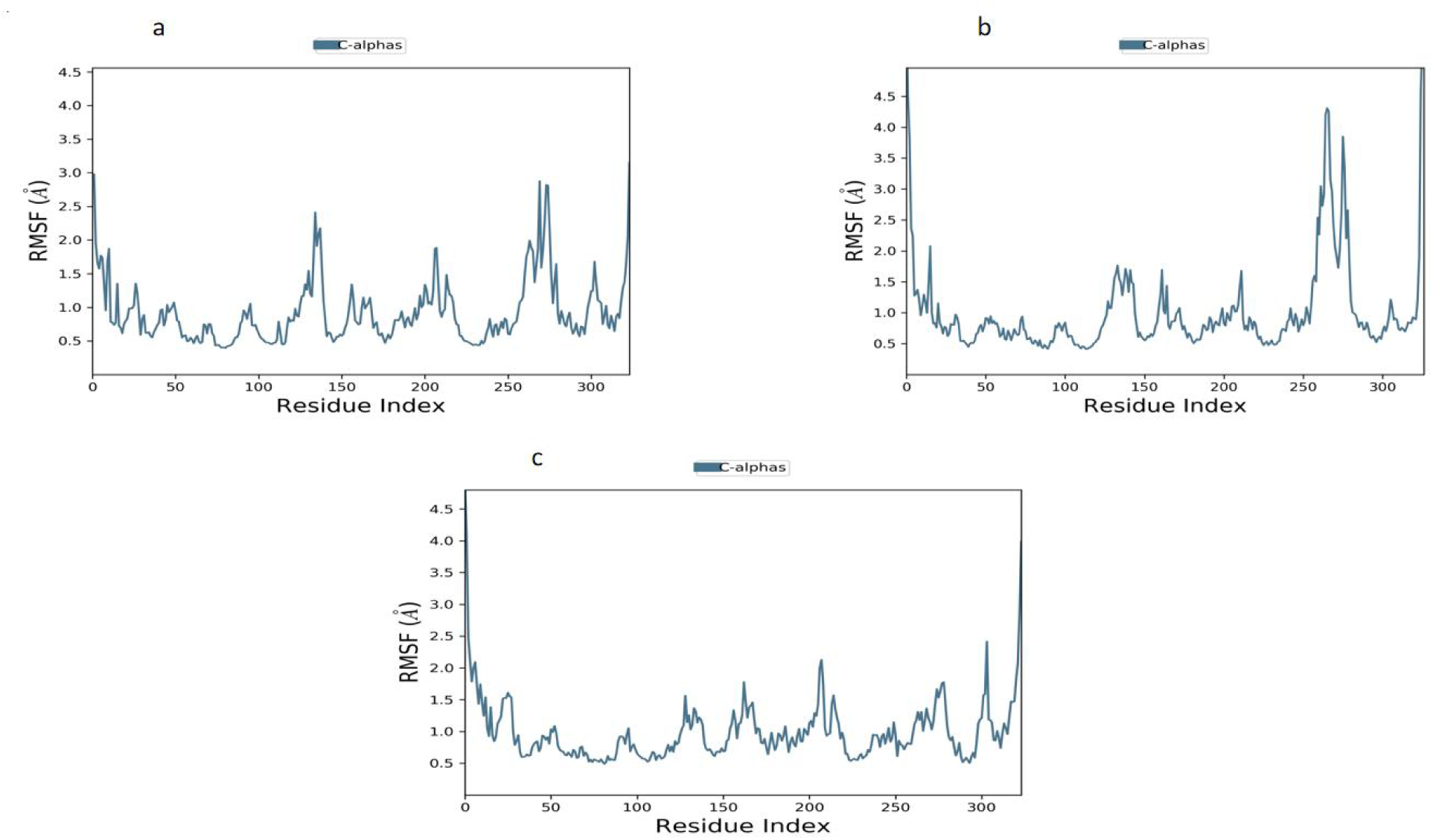
(a-c). 100 ns MD Simulation RMSF plots. a. Cα RMSF profile of 4KP6 during the 100 ns MD simulation with LTS0048837; b. Cα RMSF profile of 3G4L during the 100 ns MD simulation with LTS0048837; c. Cα RMSF profile of 4KP6 during the 100 ns MD simulation with roflumilast.

#### 3.5.3. Protein-Ligand Contacts and Interaction Persistence

Protein-ligand contact histograms (Figure 16(a-c)) were used to compare interaction persistence across the simulated complexes. In the 4KP6-LTS0048837 complex, the most persistent contacts involved ASP392, GLN443 and TYR233, with ASP392 showing a dominant hydrogen-bond contribution and GLN443 showing hydrogen-bond/water-bridge contributions. TYR233 showed hydrogen-bond/water-bridge and hydrophobic contact contributions. TYR403 and PHE446 also contributed meaningfully to the interaction network. The total contact fraction for ASP392 and GLN443 approached ∼1.8 and ∼1.35, respectively, indicating that more than one contact type occurs with the same residue during parts of the simulation. In the 3G4L-LTS0048837 complex, strong contacts were observed with ASP484 and TYR325; the interaction profile was also distributed across residues such as HIS326, MET439, TYR495, GLN535, and PHE538. For the 4KP6-roflumilast complex, the highest contact contributions involved GLN443, PHE446, ASP392, PHE414, ILE410, and TYR233.

**Figure 16.**
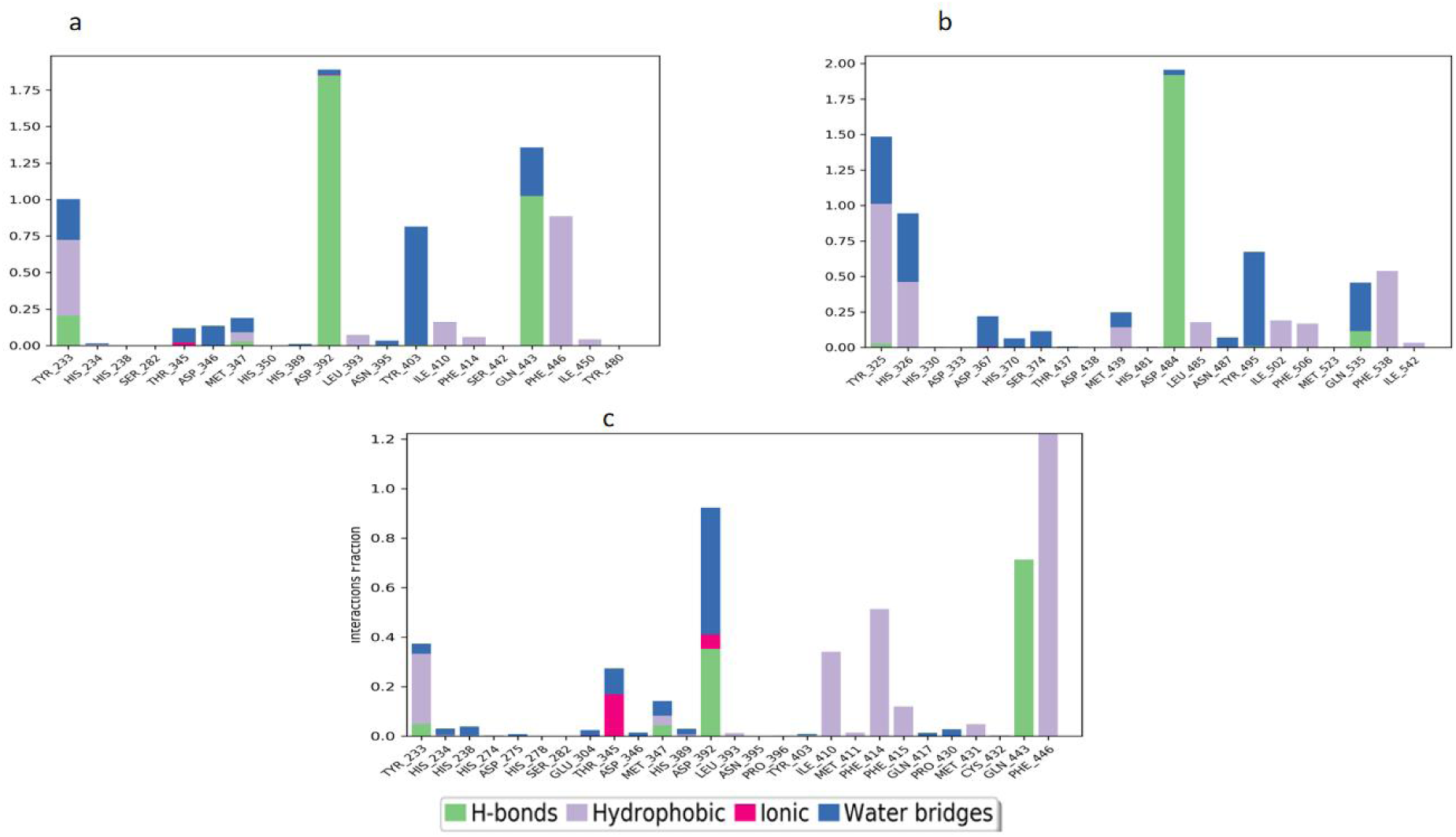
(a-c). 100 ns Protein-Ligand contact histograms. a. Protein-ligand contact histogram for the 4KP6-LTS0048837 complex during the 100 ns MD simulation; b. protein-ligand contact histogram for the 3G4L-LTS0048837 complex during the 100 ns MD simulation; c. protein-ligand contact histogram for the 4KP6-roflumilast complex during the 100 ns MD simulation.

**Figure 17.**
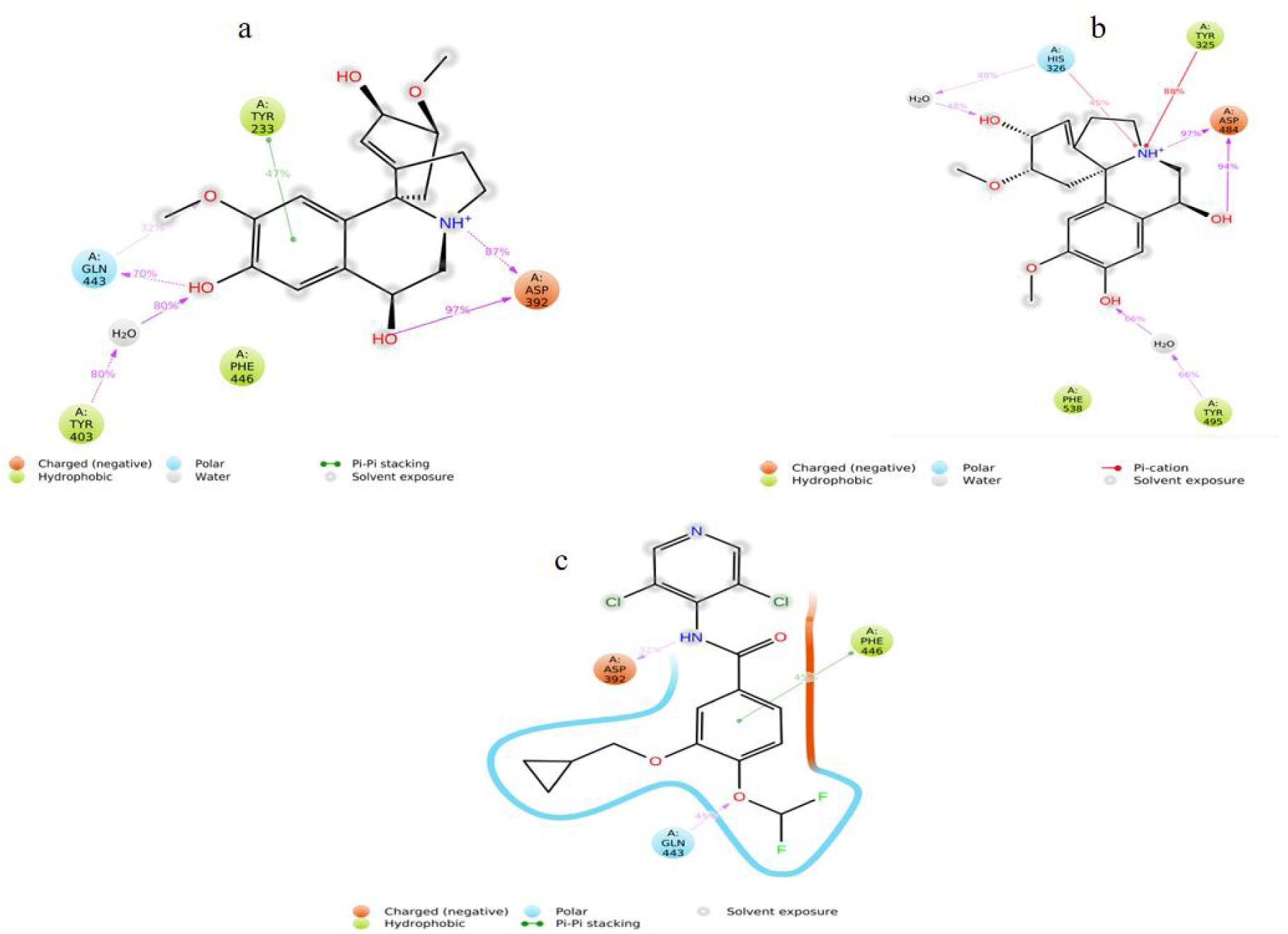
(a-c). Dynamic 2D ligand-protein contact maps from the 100 ns MD simulations. **a.** 2D ligand-protein contact map for the 4KP6-LTS0048837 complex. Contacts shown are those occurring for more than 30% of the simulation time; **b.** 2D ligand-protein contact map for the 3G4L-LTS0048837 complex. Contacts shown are those occurring for more than 30% of the simulation time; **c.** 2D ligand-protein contact map for the 4KP6-roflumilast complex. Contacts shown are those occurring for more than 30% of the simulation time

The 2D ligand-protein contact maps and contact timeline plots (Figures 17 and 18) complement the histogram-based analysis by showing both the percentage persistence of individual residue contacts and their temporal distribution across the trajectory. Only ligand-protein contact maps with greater than 30% percentage persistence are shown. In the 4KP6-LTS0048837 complex, the 2D contact map retained ASP392 as the dominant charged contact, with contacts of approximately 97% and 87%, while GLN443 contributed a persistent polar/water-mediated interaction network of approximately 70-80%. TYR403 also participated through a water-mediated contact of approximately 80%, and TYR233 contributed a pi-pi stacking contact of approximately 47%. The corresponding timeline plot showed sustained, recurrent contacts with TYR233, ASP392, TYR403, GLN443, and PHE446 across most of the 100 ns trajectory, supporting persistent binding-site engagement. In the 3G4L-LTS0048837 complex, the 2D contact map highlighted ASP484 as the dominant charged contact, with approximately 97% and 94% contact persistence, together with TYR325 (88%), TYR495-water (66%), HIS326-water (48%), and PHE538. For the 4KP6-roflumilast complex, the 2D map showed fewer contacts above the 30% reporting threshold, mainly GLN443 (45%), PHE446 (45%), and ASP392 (32%).

**Figure 18.**
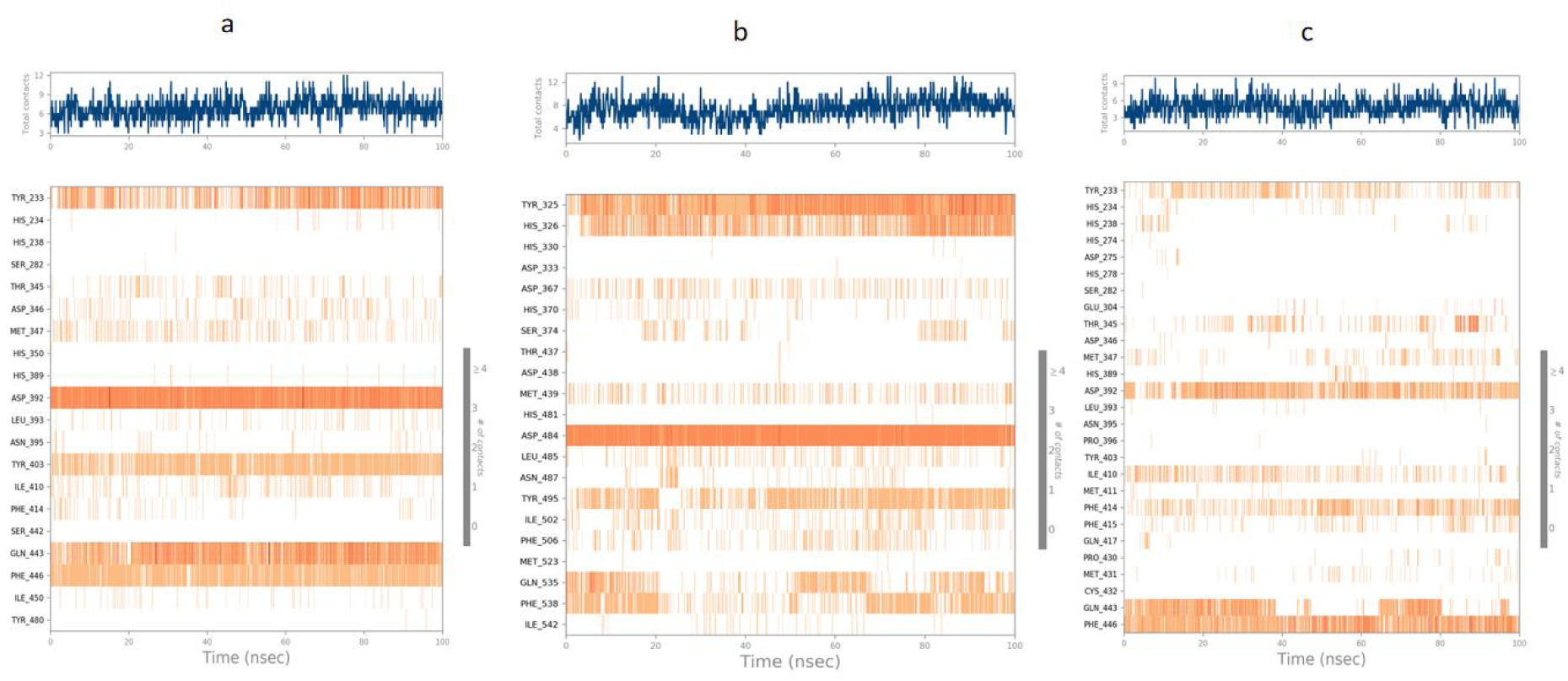
(a-c). Timeline representation of protein-ligand contacts during the 100 ns MD simulations. a. Contact timeline plot for the 4KP6-LTS0048837 complex, showing the total number of contacts over time and the residue-level temporal persistence of contacts; b. Contact timeline plot for the 3G4L-LTS0048837 complex, showing the total number of contacts over time and the residue-level temporal persistence of contacts; c. Contact timeline plot for the 4KP6-roflumilast complex, showing the total number of contacts over time and the residue-level temporal persistence of contacts.

### 3.6. ADMET and Physicochemical Properties

The physicochemical and ADMET properties of the four lead compounds and roflumilast are shown in Table 3. The four leads that were selected met Lipinski’s Rule-of-Five without a single violation. Molecular weights ranged from 238.67 to 342.39 Da, while calculated iLogP values ranged from 1.25 to 3.13. TPSA values ranged from 51.16 to 82.39 Å². All selected compounds showed high predicted gastrointestinal absorption.

**Table 3.**
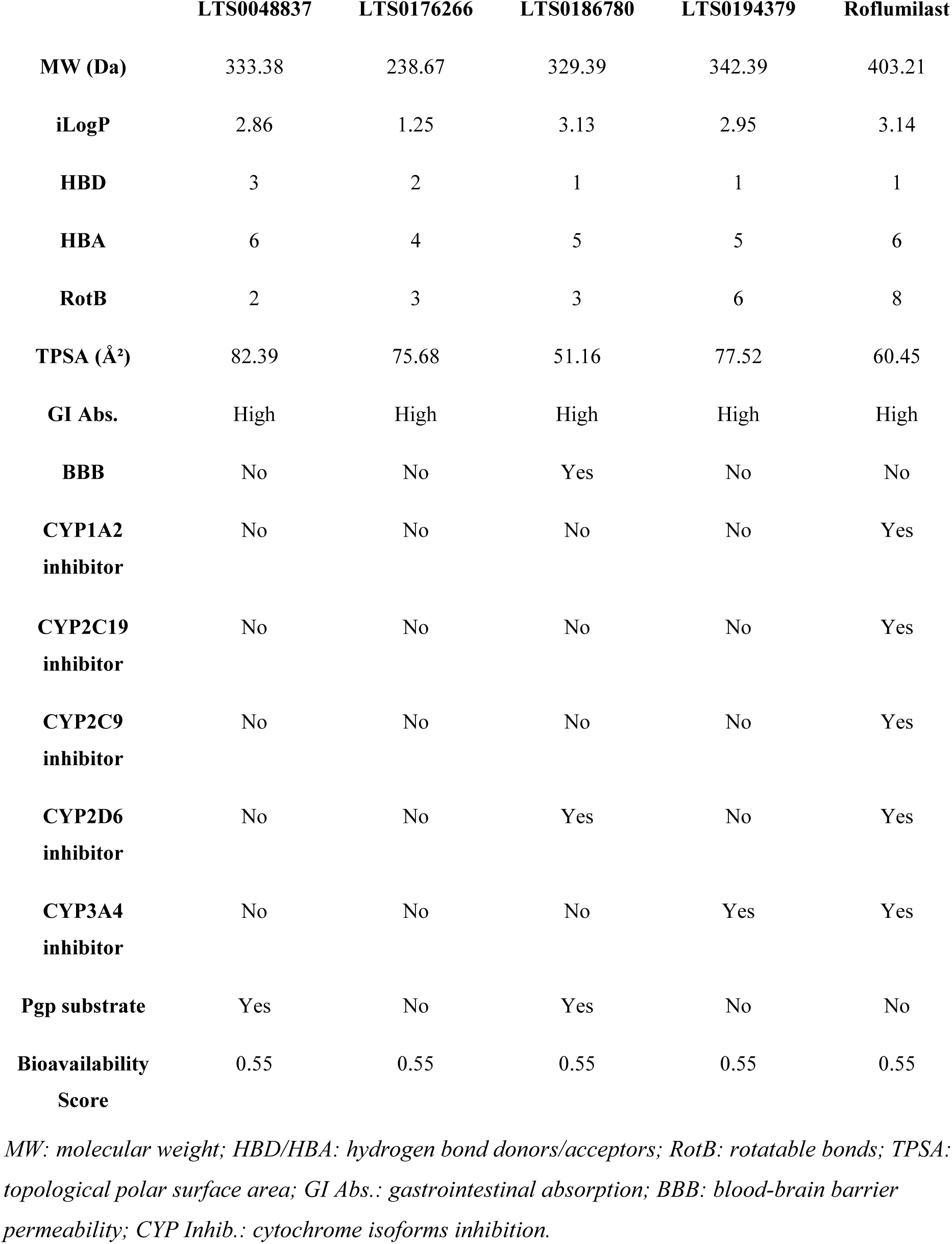
Physicochemical and ADMET properties of top four lead compounds and roflumilast.

BBB permeability prediction showed that LTS0048837, LTS0176266, and LTS0194379 were not predicted to cross the blood-brain barrier, while LTS0186780 was predicted to be BBB-permeant. Regarding P-glycoprotein (P-gp) substrate status, LTS0048837 and LTS0186780 were predicted to be P-gp substrates, whereas LTS0176266 and LTS0194379 were not. CYP inhibition profiling showed that LTS0048837 and LTS0176266 were not predicted to inhibit any of the evaluated CYP isoforms. LTS0186780 was predicted to inhibit CYP2D6 only, while LTS0194379 was predicted to inhibit CYP3A4 only. In contrast, roflumilast was predicted to inhibit all evaluated CYP isoforms, including CYP1A2, CYP2C19, CYP2C9, CYP2D6, and CYP3A4. All selected leads had a bioavailability score of 0.55, similar to roflumilast.

## 4. Discussion

In the two-stage computational strategy reported in this study, selective inhibitors of PDE4B were discovered from natural products, yielding LTS0048837 as the lead candidate with superior predicted potency and selectivity over roflumilast. The convergent evidence from ML bioactivity prediction, hierarchical docking, binding-pocket comparison, post-docking MM-GBSA binding free-energy calculations, and 100 ns MD simulations establishes a multi-method computational case for PDE4B-selective binding, a central pharmacological challenge in this target space.

The clinical motivation for PDE4B-selective inhibition is well established: PDE4D inhibition drives the principal tolerability liabilities of roflumilast (nausea, diarrhea, weight loss) through emetic pathway modulation, while PDE4B inhibition is the primary mediator of anti-inflammatory efficacy in pulmonary tissue [12–15]. The computational selectivity profile suggests that LTS0048837 preferentially engages PDE4B over PDE4D, with structural analysis identifying a key non-conserved active-site residue, Tyr233, as a plausible contributor to selectivity.

The ML pipeline achieved an AUC-ROC of 0.948 and MCC of 0.785; the MCC is of particular significance as it provides a balanced assessment under potential class imbalance and is increasingly recommended as the primary metric for binary classification [35].

The natural products used as the screening library were deliberately chosen from the LOTUS database. Natural products possess evolved three-dimensional molecular architectures and conformational complexity often absent from synthetic compound libraries, and have historically been disproportionately productive as lead sources for enzyme inhibitors [18]. LTS0048837 and the other lead compounds have not been previously reported as a PDE4B inhibitor to the best of our knowledge, representing structurally novel natural product scaffolds for this target and underscoring the potential of the natural product chemical space for PDE4 isoform-selective lead discovery.

In addition to the better docking and post-docking MM-GBSA results of PDE4B-LTS0048837 complex, compared to the PDE4D-LTS0048837 complex, the MD simulation results add an important dynamic layer . In the PDE4B-LTS0048837 complex, protein RMSD stabilized after the early equilibration phase and the ligand RMSD remained within a narrow range, indicating that the docked pose was stable throughout the 100 ns trajectory. The differential behavior observed in the PDE4D-LTS0048837 simulation provides a good context for the selectivity hypothesis.

Although the ligand was still located in the binding pocket of the PDE4D, the PDE4D protein backbone showed a more progressive RMSD increase and the RMSF profile contained more pronounced localized peaks than the PDE4B complex. This suggests that the PDE4D pocket may require greater conformational adjustment to accommodate LTS0048837, whereas the PDE4B pocket supports a more stable binding arrangement.

The contact histogram further showed that LTS0048837 maintained persistent interactions with key PDE4B binding-site residues, especially ASP392, GLN443 and TYR233, while also engaging residues such as TYR403, PHE446, and ILE410.

The 2D contact maps and timeline plots strengthen this interpretation by demonstrating that the key contacts were not isolated events but recurrent interactions distributed across the trajectory. In particular, the PDE4B-LTS0048837 complex maintained repeated contacts with ASP392, GLN443, TYR403, TYR233, and PHE446, whereas roflumilast showed fewer high-persistence contacts above the 30% threshold in the corresponding 2D contact map.

It is worth noting that of the top four compounds, LTS0194379 did not demonstrate superior PDE4B docking score selectivity. However, it showed a better PDE4B MMGBSA score. Medicinal chemistry optimization targeting PDE4B-discriminating residues could potentially enhance the scaffold.

The ADMET predictions further support the drug-like potential of the selected lead compounds. Their molecular weights, iLogP values, TPSA values, and absence of Lipinski violations suggest acceptable physicochemical properties for small-molecule development [24,31]. The high predicted gastrointestinal absorption observed for all four leads is also favourable, particularly for oral delivery consideration.

The BBB prediction is relevant because the intended therapeutic context is COPD, where the primary pharmacological site of action is peripheral lung tissue rather than the central nervous system. Therefore, the absence of predicted BBB penetration for LTS0048837, LTS0176266, and LTS0194379 may be advantageous by reducing the likelihood of unwanted CNS exposure. However, LTS0186780 was predicted to be BBB-permeant, which may require closer safety consideration during future optimization.

The P-glycoprotein substrate prediction suggests that LTS0048837 and LTS0186780 may be more susceptible to transporter-mediated efflux, which could influence tissue distribution, intracellular exposure, or oral bioavailability, although this requires experimental validation [34]. CYP inhibition profiling also suggests that the lead compounds may have a lower predicted drug-drug interaction burden than roflumilast. Notably, LTS0048837 and LTS0176266 showed no predicted inhibition of the evaluated CYP isoforms, whereas roflumilast was predicted to inhibit all five CYP isoforms assessed. Overall, the ADMET profiles of the selected leads compare favorably with roflumilast, particularly in terms of reduced predicted CYP inhibition burden, favorable gastrointestinal absorption, acceptable physicochemical properties, and generally limited BBB penetration.

Some limitations of this study warrant acknowledgment. All findings are computational predictions and require experimental validation, including enzymatic IC50 determination against recombinant PDE4B and PDE4D, cellular cAMP elevation assays, and in vivo pharmacokinetic profiling before a strong clinical relevance can be established. The MD simulations were limited to 100 ns and may not capture all slow conformational transitions; extended simulations, replicate trajectories, enhanced sampling, or trajectory-based free-energy approaches could provide additional confidence in stability conclusions. Furthermore, the selectivity assessment relied on rigid-receptor docking and post-docking single-pose MM-GBSA, which do not fully account for induced-fit conformational changes or full thermodynamic ensemble effects that may differentially affect PDE4B and PDE4D binding.

## 5. Conclusions

This study introduced a computational workflow that combines SHAP-interpretable machine learning and structure-based drug design for the discovery of PDE4B selective inhibitors from the LOTUS database of natural products. The ML model enabled an efficient bioactivity-driven prioritization of the 276,518-compound LOTUS library. Multi-stage structure-based evaluation, hierarchical docking, post-docking MM-GBSA, isoform cross-docking, binding-pocket comparison, and 100 ns MD simulation, identified LTS0048837 as the top lead candidate, demonstrating superior predicted potency to roflumilast, stable retention in the PDE4B active site, and, consequently, a stronger overall PDE4B-selectivity profile than the PDE4D complex. SHAP analysis allowed interpretability by identifying the ligand features driving the machine-learning predictions. The favorable ADMET profile of the lead candidates further supports their drug-like potential. These findings constitute a strong computational foundation warranting experimental follow-up, including enzymatic inhibition assays, PDE4B/PDE4D selectivity assays, cellular anti-inflammatory assays, and in vivo COPD model evaluation, toward the development of safer PDE4B-selective therapeutics with an improved tolerability profile over existing non-selective PDE4 inhibitors.

## Notes

### Competing Interest Statement

The authors have declared no competing interest.

